# PURE: Policy-guided Unbiased REpresentations for structure-constrained molecular generation

**DOI:** 10.1101/2025.05.21.655002

**Authors:** Abhor Gupta, Barathi Lenin, Sean Current, Rohit Batra, Balaraman Ravindran, Karthik Raman, Srinivasan Parthasarathy

## Abstract

Structure-constrained molecular generation (SCMG) generates novel molecules that are structurally similar to a given molecule and have optimized properties. Deep learning solutions for SCMG are limited in that they are pre-disposed towards existing knowledge, and they suffer from a natural impedance mismatch problem due to the discrete nature of molecules, while deep learning methods for SCMG often operate in continuous space. Moreover, many task-specific evaluation metrics used during training often bias the model towards a particular metric -”metric-leakage”. To overcome these shortcomings, we propose Policy-guided Unbiased REpresentations (PURE) for SCMG that learn within a framework simulating molecular transformations for drug synthesis. PURE combines self-supervised learning with a policy-based reinforcement-learning (RL) framework, thereby avoiding the need for external molecular metrics while learning high-quality representations that incorporate an inherent notion of similarity specific to the given task. Along with a semi-supervised training design, PURE utilizes template-based molecular simulations to better explore and navigate the discrete molecular search space. Despite the lack of metric biases, PURE achieves competitive or superior performance than state-of-the-art methods on multiple benchmarks. Our study emphasizes the importance of reevaluating current approaches for SCMG and developing strategies that naturally align with the problem. Finally, we illustrate how our methodology can be applied to combat drug resistance, by identifying sorafenib-like compounds as a case study.

## 1 Introduction

The discovery and development of novel drugs and functional materials are vital in modern science. However, navigating the vast chemical space in traditional *in silico* drug discovery still presents a significant challenge. Structure-Constrained Molecular Generation (SCMG) offers a powerful approach to address this limitation. SCMG is the technique of using existing molecules to guide the creation of new ones with similar structures and potentially improved properties. This technique harnesses the knowledge of existing successful molecules to guide the generation of new compounds with similar core structures. SCMG enables researchers to explore targeted regions within the chemical space by focusing on specific chemical frameworks. SCMG finds further utility throughout the drug discovery process, specifically aiding in lead optimization and the identification of novel therapeutics to overcome resistance developed by patients or bacteria [1–3].

With recent advances in deep learning, much attention has been put into developing methods for drug discovery and, consequently, towards SCMG. Several deep learning solutions have been proposed for SCMG, predominantly employing generative methods and reinforcement learning (RL). You et al. [4] used RL to generate molecular graphs. They utilized property rewards along with an adversarial loss to affix the distribution of their generated molecules similar to the set of target molecules. Zhou et al. [5] trained a Deep Q-Network (DQN) to generate molecular graphs similar to You et al. [4] without pretraining to allow for broader exploration. Işık and Tan [6] build upon Zhou et al. [5] and used graph networks for molecular representations. Jeon and Kim [7] also builds upon Zhou et al. [5] while giving rewards for a docking score along with a property score. Ståhl et al. [8] utilized BiLSTM-based actor-critic architectures to replace fragments in molecules to improve chemical properties. They encoded fragments in a binary string using a binary tree where similar fragments are close to each other, further allowing them to control the similarity of molecules generated. Olivecrona et al. [9] used RL to fine-tune an RNN to generate SMILES strings for molecules with desired properties. Tan et al. [10] trained a transformer over SMILES and fine-tuned using a reward combining SC score and a thresholded Tanimoto similarity. Jin et al. [11] proposed an encoder-decoder architecture to generate a tree representing the molecule’s substructure and then combine it into a molecule using a message-passing network. Jin et al. [12] built upon their work by adding an attention mechanism to the decoder and integrating an adversarial component to the training. Fu et al. [13] built upon Jin et al. [12]’s work using a copy and refine strategy, where at each step, it selects whether to copy the following substructure from the input or to generate a new substructure. Jin et al. [14] proposed a hierarchical encoder-decoder architecture to generate molecular graphs using structural motifs as building blocks. Fan et al. [15] enhanced their work by suggesting a semi-supervised back-translation method for generating synthetic data to augment their training. Barshatski and Radinsky [16] proposed an unsupervised approach to generate molecular SMILES that uses a double-cycle training scheme for molecular translation. Choi et al. [3] used an objective with contractive and margin losses to learn structural similarity and fine-tune for property optimization using RL.

Several other works exist in the literature, including sampling-based methods [17], search-based methods [18–20], auto encoders and VAEs [21–24], flow models [24–28], GANs [29], transformers [30], diffusion ([31] and genetic algorithms [32]. However, navigating the vast chemical space in traditional *in silico* drug discovery still presents a significant challenge. While many methods described above can navigate the vast chemical space to identify molecules that satisfy chemical rules, a considerable challenge remains that in only a small percentage of these molecules are practically viable - synthesizable and pharmacologically relevant. Furthermore, the space of molecules is discrete, whereas generative methods in several proposed methods operate in continuous space. Additionally, many of these methods require high-quality training data, which is difficult to obtain and leverage for existing SCMG methods. Another significant issue with such methods is their exposure to evaluation metrics during training. This “metric leakage” can cause overfitting and bias in the models towards exploiting weaknesses in the metrics. For example, optimizing for octanol-water partition coefficient (logP) can lead deep learning methods to increase molecule size, a known artifact of the metric [33], rather than achieving genuine improvements in the desired property. This issue is further exacerbated because the computation of most molecular metrics only partially captures the molecular properties in question [34].

To address these fundamental challenges, we take a step back from the recent trend of research towards SCMG to design a self-supervised and metric-agnostic framework for this task. Towards this effect, we propose PURE: Policy-guided Unbiased REpresentations for SCMG. Along with a semi-supervised and metric-agnostic training design, PURE utilizes template-based molecular simulations, which improve synthesizability and constrain the search space to be discrete while also employing the exploratory capabilities of RL to navigate the discrete molecular space.

The crux of our method is the observation that approximation under source-to-target path prediction in a similarity learning framework leads to SCMG (Figure 1 shows an overview of PURE). To avoid using similarity and property metrics during training, we first decouple similarity and property optimization into different framework phases. The similarity optimization phase focuses on the self-supervised auxiliary task of source-to-target path prediction. Training under the goal-conditioned RL paradigm allows PURE to learn high-quality molecular representations. The inner product between the learned embeddings can be used as a similarity score [35]. In the generation phase, this learned notion of similarity is used with beam search to generate molecules similar to a target molecule. This leads to many unbiased alternative candidates for the target since neither the training nor generation phases are exposed to any of the molecular metrics. Finally, among the generated molecules, the molecules with the desired properties are selected by curating the top *k* molecules based on the required combined objective from the set of generated molecules. We believe our work suggests a new perspective from recent efforts and excites the research community to broaden their exploration for developing new and effective methods for drug discovery.

**Fig. 1.**
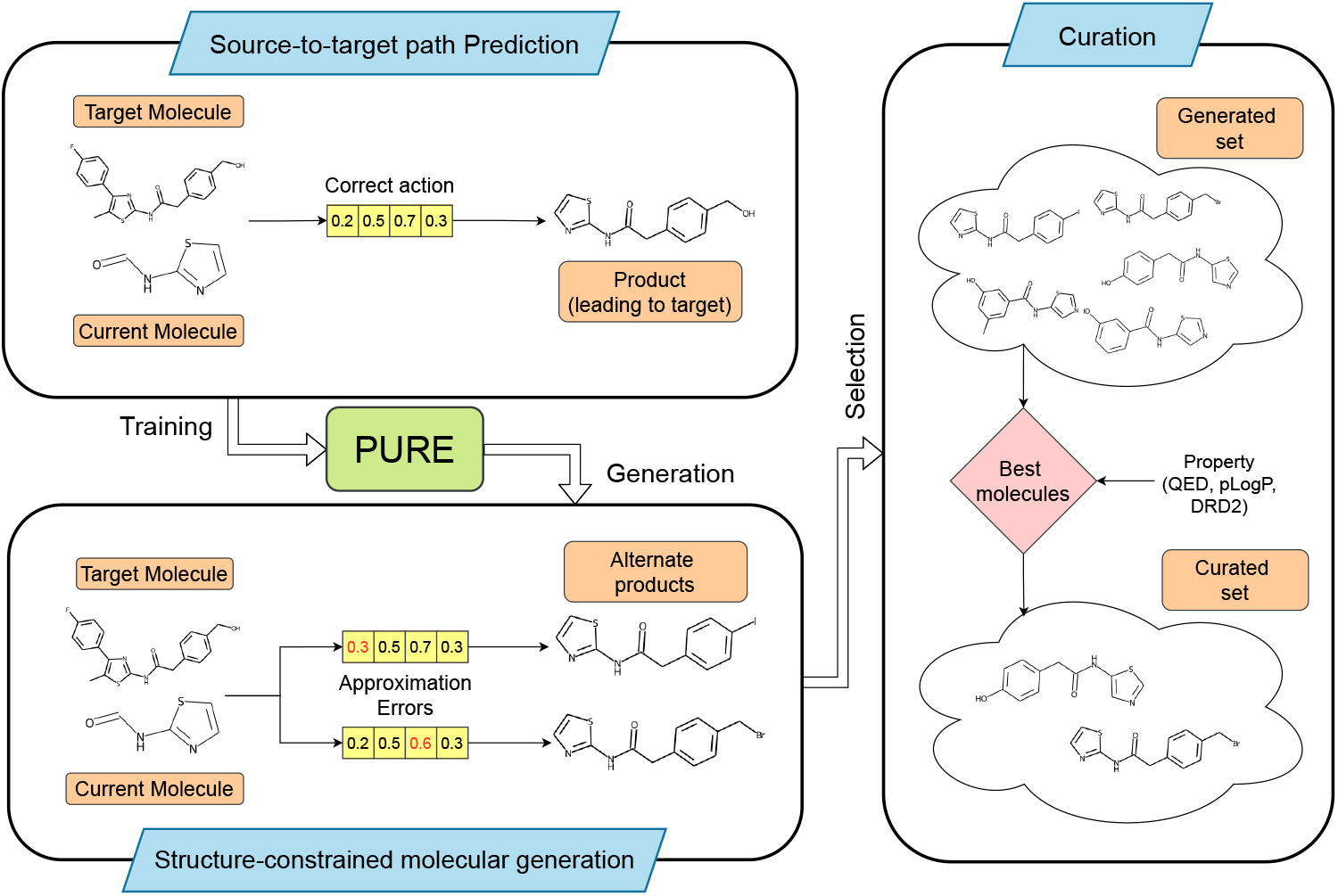
Overview of PURE. PURE is trained on the task of **source-to-target path prediction** to predict the correct actions given the current molecule and target molecule. Learning under a similarity learning framework allows zero-shot transfer of PURE on **SCMG** such that approximation errors lead to generating similar molecules. Lastly, a **curation** step is applied that selects the best molecules according to the desired properties or their combination, resulting in the final curated set of optimized molecules.

We compare results for SCMG on four benchmarks: QED, DRD2, pLogP04, and pLogP06 from Choi et al. [3]. Despite not being exposed to similarity or property metrics during training or generation, we show that PURE achieves comparable or better performance than previous methods. Unlike previous methods that require training a different model for each similarity–property combination, **a *single* model trained under the PURE framework can generate optimized molecules for each property benchmark**. Furthermore, we conduct a case study focusing on Sorafenib, an anti-cancer drug. Our results demonstrate that PURE successfully generates a significantly larger number of molecules compared to COMA[3] while exhibiting improved properties and fewer violations. This case study highlights the versatility and efficacy of the PURE framework in generating drug-like molecules with desired characteristics.

Furthermore, to ensure the validity of molecules, we follow a fragment replacement strategy inspired by [10], but instead of replacing fragments or “reaction rules” on the lead candidate, we operate on smaller precursor molecules, thereby simulating a stepwise drug synthesis process. These reaction rules, mined and extracted from USPTO-MIT [36], induce synthesizability into the design of the molecules and also provide a possible path of synthesis as part of the generation process, which is another advantage of PURE over other existing generative SCMG models.

## 2 Results

In this section, we report how PURE consistently outperforms existing methods across multiple popular benchmarks for SCMG, demonstrating superior diversity, novelty, and synthesizability. Furthermore, we demonstrate the effectiveness of PURE via a case study aimed at generating alternative drug candidates to overcome the challenge of sorafenib resistance.

### 2.1 Benchmarking PURE

We report our results for benchmarks QED, DRD2, LogP04, and LogP06 in Tables 1, 2, 3, and 4, respectively. We show comparisons against the baseline models on the metrics validity, property, improvement, similarity, novelty, and diversity for the four benchmark models. We compare with COMA [3] and the 7 baselines used in their work: JTVAE [11], VJTNN, VJTNN+GAN [12], CORE [13], HierG2G [14], HierG2G+BT [15], and UGMMT [16]. During our generation process, our model generates about 2,000 candidate molecules for each source-target pair, which are then filtered by selecting the top 20 molecules for each target by some linear combination of similarity and property scores: (*x*.*similarity* + *y*.*property*). For QED and DRD2 benchmarks, we show results for (*x, y*) ∈ {(0, 1), (1, 0), (1, 1)} while for the logP benchmarks, we show results for (*x, y*) ∈ {(0, 1), (1, 0), (25, 1)} to accommodate for the range of logP values. Further details of benchmarks, metrics, baselines and evaluation are present in Section 4.8.

**Table 1.**
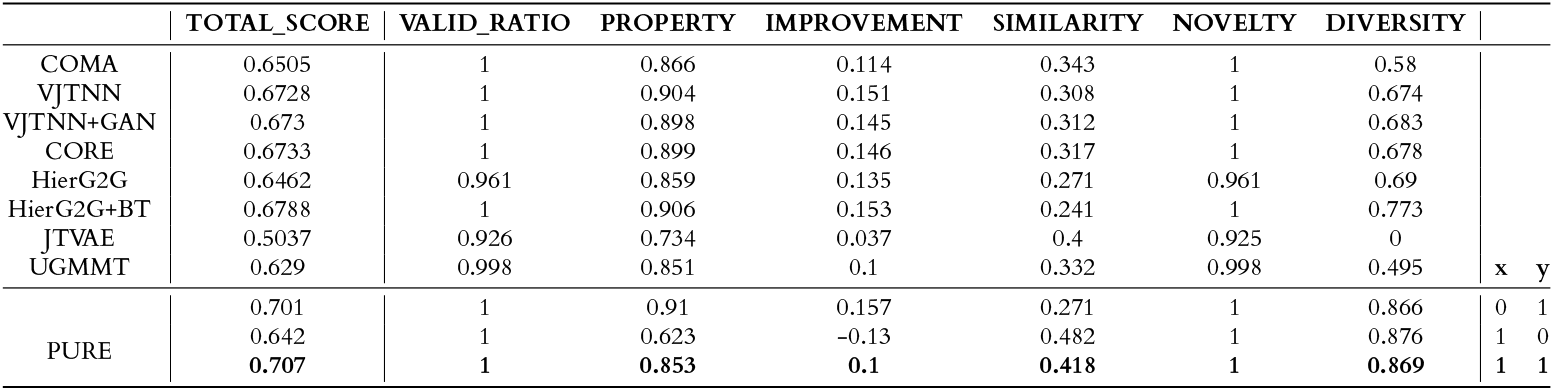
Comparison on QED benchmark.

**Table 2.** Comparison on DRD2 benchmark.

|  | TOTAL_SCORE | VALID_RATIO | PROPERTY | IMPROVEMENT | SIMILARITY | NOVELTY | DIVERSITY |  |
| --- | --- | --- | --- | --- | --- | --- | --- | --- |
| COMA | 0.7142 | 1 | 0.799 | 0.792 | 0.329 | 0.999 | 0.366 |  |
| VJTNN | 0.6787 | 0.999 | 0.803 | 0.797 | 0.344 | 0.696 | 0.433 |  |
| VJTNN+GAN | 0.7137 | 0.999 | 0.787 | 0.78 | 0.324 | 0.885 | 0.507 |  |
| CORE | 0.6947 | 0.999 | 0.8 | 0.793 | 0.338 | 0.779 | 0.459 |  |
| HierG2G | 0.6922 | 0.981 | 0.68 | 0.673 | 0.258 | 0.971 | 0.59 |  |
| HierG2G+BT | 0.6783 | 1 | 0.808 | 0.802 | 0.336 | 0.671 | 0.453 |  |
| JTVAE | 0.4077 | 0.935 | 0.091 | 0.084 | 0.402 | 0.934 | 0 |  |
| UGMMT | 0.7088 | 0.999 | 0.737 | 0.73 | 0.261 | 0.996 | 0.53 | x y |
| PURE | 0.772 | 1 | 0.761 | 0.755 | 0.27 | 1 | 0.846 | 0 1 |
|  | 0.576 | 1 | 0.029 | 0.022 | 0.555 | 1 | 0.851 | 1 0 |
|  | **0.771** | **1** | **0.73** | **0.723** | **0.329** | **1** | **0.847** | **1 1** |

**Table 3.** Comparison on LogP04 benchmark.

|  | TOTAL_SCORE | VALID_RATIO | PROPERTY | IMPROVEMENT | SIMILARITY | NOVELTY | DIVERSITY |  |
| --- | --- | --- | --- | --- | --- | --- | --- | --- |
| COMA | 1.5883 | 0.995 | 1.945 | 4.663 | 0.308 | 0.995 | 0.624 |  |
| VJTNN | 1.4113 | 0.975 | 1.489 | 4.095 | 0.316 | 0.975 | 0.618 |  |
| VJTNN+GAN | 1.4497 | 0.99 | 1.543 | 4.242 | 0.326 | 0.99 | 0.607 |  |
| CORE | 1.4063 | 0.984 | 1.431 | 4.087 | 0.324 | 0.984 | 0.628 |  |
| HierG2G | 1.3952 | 0.948 | 1.51 | 4.052 | 0.266 | 0.948 | 0.647 |  |
| HierG2G+BT | 1.3735 | 0.981 | 1.335 | 3.983 | 0.269 | 0.981 | 0.692 | x y |
| PURE | 3.075 | 1 | 6.373 | 9.118 | 0.112 | 1 | 0.849 | 0 1 |
|  | 0.988 | 1 | -0.052 | 2.692 | 0.404 | 1 | 0.884 | 1 0 |
|  | **1.993** | **1** | **3.008** | **5.752** | **0.324** | **1** | **0.874** | **25 1** |

**Table 4.** Comparison on LogP06 benchmark.

|  | TOTAL_SCORE | VALID_RATIO | PROPERTY | IMPROVEMENT | SIMILARITY | NOVELTY | DIVERSITY |  |
| --- | --- | --- | --- | --- | --- | --- | --- | --- |
| COMA | 1.3285 | 0.958 | 1.313 | 3.86 | 0.383 | 0.958 | 0.499 |  |
| VJTNN | 0.9812 | 0.826 | 0.768 | 2.772 | 0.399 | 0.825 | 0.297 |  |
| VJTNN+GAN | 1.128 | 0.881 | 0.987 | 3.222 | 0.397 | 0.881 | 0.4 |  |
| CORE | 1.0617 | 0.859 | 0.871 | 3.037 | 0.396 | 0.859 | 0.348 |  |
| HierG2G | 0.9943 | 0.825 | 0.788 | 2.828 | 0.344 | 0.825 | 0.356 |  |
| HierG2G+BT | 1.0285 | 0.849 | 0.83 | 2.908 | 0.374 | 0.849 | 0.361 | x y |
| PURE | 3.075 | 1 | 6.373 | 9.118 | 0.112 | 1 | 0.849 | 0 1 |
|  | 1.372 | 1 | 1.112 | 3.856 | 0.384 | 1 | 0.881 | 1 0 |
|  | **2.012** | **1** | **3.068** | **5.812** | **0.32** | **1** | **0.873** | **25 1** |

For most benchmarks, PURE outperforms previous state-of-the-art methods across almost all metrics. For the QED benchmark, PURE outperforms the other methods on valid_ratio, similarity, novelty and diversity for (*x* = 1, *y* = 1) and (*x* = 1, *y* = 0). For (*x* = 0, *y* = 1), PURE achieves the best property scores while suffering on similarity and retaining the same level of performance on other metrics. Similar trends are observed in the other benchmarks as well. PURE lags behind DRD2 in the property scores, while the opposite is true for LogP benchmarks, where the property scores are significantly higher than the other methods. In fact, for LogP04, PURE comes close to outperforming the other methods on every single benchmark.

As seen in the Tables 1-4, the coefficients *x* and *y* are critical in selecting molecules with the desired properties. We plot the relationship between the coefficients and the corresponding molecular metrics in Fig 2, for the generated set of molecules. Earlier works often utilize an equal weighting between similarity and property metrics (e.g., *x* = 1, *y* = 1). However, due to the interdependence and disparate scales of these metrics, a equal weighting scheme falls short of effectively encapsulating the desired molecular characteristics. In fact, optimal weightings for these metrics are often task-specific and difficult to predetermine. PURE addresses these challenges by decoupling coefficient selection from the generation process, enabling flexible post-generation tuning. This approach significantly reduces computational costs compared to training separate models for different coefficient combinations. Ultimately, PURE helps chemists overcome the hurdle of specifying exact property requirements for drug candidates prior to model training.

**Fig. 2.**
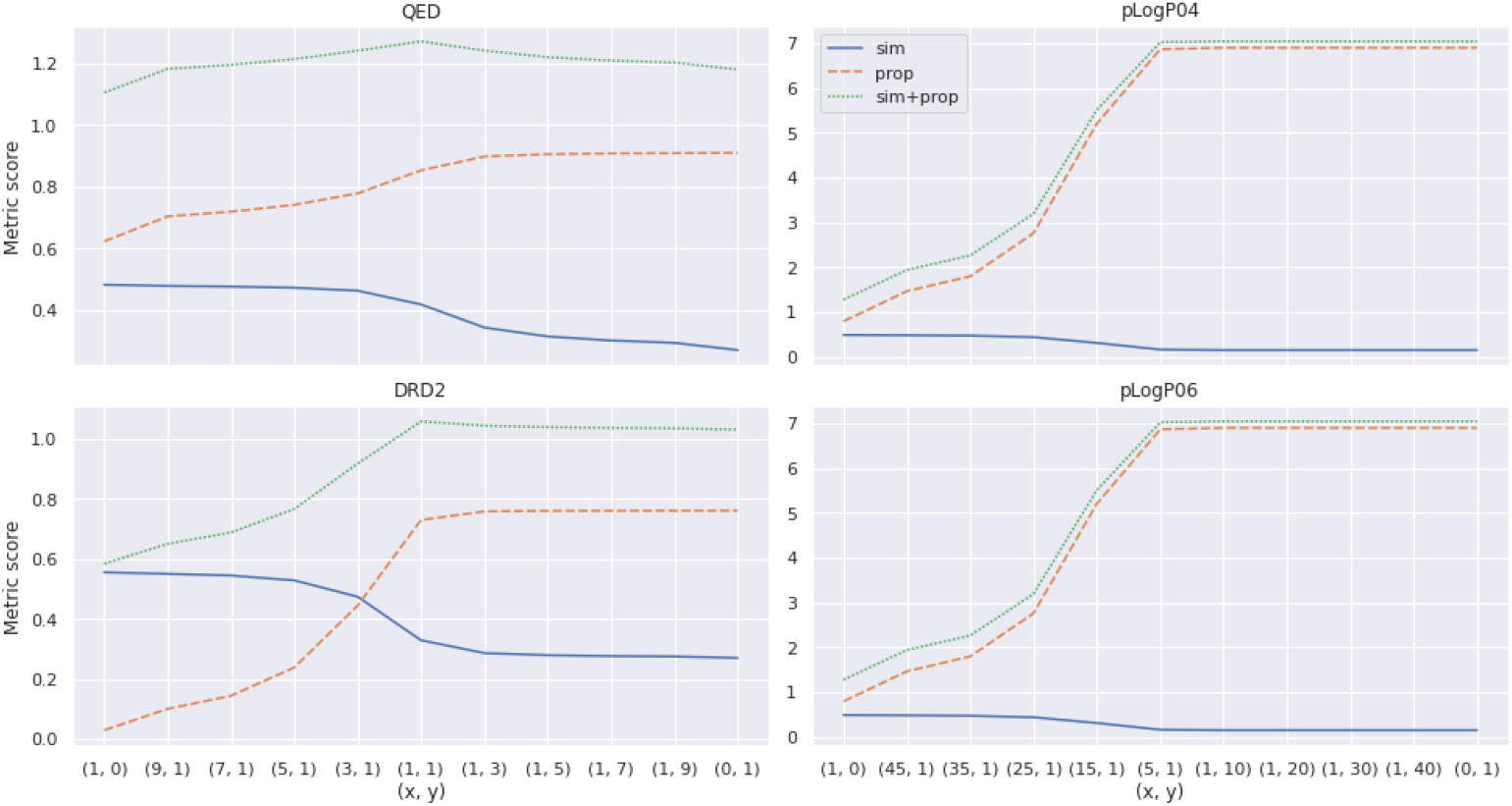
The plots show the effect of parameters *x* and *y* on the desired property metrics on the generated set of molecules for each benchmark; ‘sim’ represents Tanimoto similarity, ‘prop’ represents the desired property for the benchmark (QED, DRD2 or pLogP) and ‘sim+prop’ shows the linear sum of the two.

### 2.2 Case Study: Addressing Sorafenib Resistance

Sorafenib is a multi-targeted kinase inhibitor approved for the treatment of advanced renal cell carcinoma, hepatocellular carcinoma and thyroid cancer [37]. Its mechanism of action involves the blocking of tumor cell proliferation and angiogenesis. Sorafenib has been shown to extend survival, but its clinical efficacy has been severely impacted by side effects and the development of drug resistance [38]. A significant mechanism contributing to this resistance is the overexpression of ABCG2, an efflux transporter that actively pumps sorafenib out of cancer cells, reducing its intracellular concentration and limiting its anti-tumor effects [39]. Several attempts have been made to circumvent such resistance, ranging from developing alternatives to sorafenib to modifying the scaffold of sorafenib to reduce its affinity towards efflux pump proteins[3, 38, 39].

This makes the drug an ideal candidate for evaluating PURE, which attempts to explore a novel chemical space and develop molecules that retain the desirable properties of the precursor while overcoming unwanted effects, such as drug resistance. COMA [3] carried out a case study where sorafenib-based molecules were evaluated against the protein ABCG2 and also one of its kinase targets, serine/threonine-protein kinase B-raf (BRAF). Similarly, we apply our model to sorafenib and evaluated its performance by comparing the results with those generated by COMA.

We follow a sequential filtration process, which yields 1,744 candidates meeting key criteria for docking simulations, as detailed in the Methods section. At each step, sorafenib and the 19 small molecules from COMA are evaluated and compared alongside the sufficiently small PURE molecules (under 500 Da). All COMA molecules also fall within this 500 Da range. Next, molecules exhibiting binding affinities greater than sorafenib towards ABCG2 are filtered, resulting in 1,134 molecules. To determine interactions with B-RAF, we analyze protein-ligand complexes using PLIP profiling. The observed interactions include hydrogen bonds, hydrophobic interactions, *π*-*π* stacking, cation-*π* interactions, halogen bonds, and salt bridges with BRAF. Sorafenib specifically forms a *π*-*π* stacking interaction with PHE 594, a halogen bond with THR 507, and several hydrophobic interactions. It also forms four unique hydrogen bonds with LYS 482, GLU 500, CYS 531, and ASP 593. A stringent threshold is applied to all PURE and COMA ligands, requiring hydrogen bonds with these same four residues in docked complexes. This filtering step excludes all COMA molecules, leaving 214 PURE molecules as lead candidates for the alternative to sorafenib.

We perform further analysis to identify the most promising drug candidates from our lead compounds. This includes ADME analysis using SwissADME to prioritize druglikeness, lead-likeness, and PAINS/Brenk filters. Depending on the stringency of these criteria, varying numbers of ligands pass the filters. Of the resulting ligands, 81 satisfy all drug-likeness filters and lack problematic PAINS and Brenk structural alerts. Further lead-likeness filtering reveals 30 PURE ligands with one or fewer violations (compared to three violations in sorafenib), which are identified as the best candidates. 15 of the “top” molecules, including 9 with no violations are shown in Table 5 (structures for some of these are shown in Figure 3(c)). It is worth noting that sorafenib does not meet certain drug-likeness criteria and has several lead-likeness issues. Therefore, we place more focus on identifying compounds that exceed these standards.

**Table 5.**
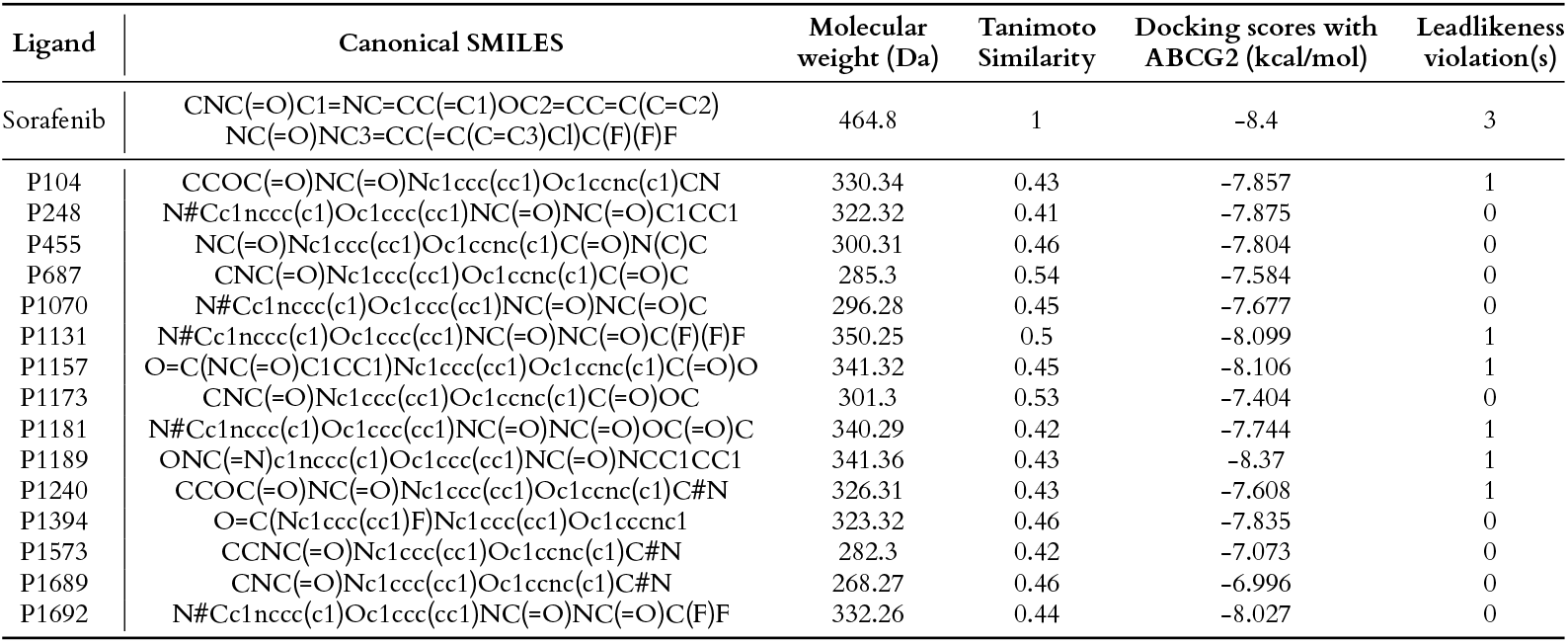
The table shows details of sorafenib, the top 9 and a few more promising molecules generated by PURE, having 1 or fewer leadlikeness violations. The molecule descriptors, docking scores against ABCG2, and Tanimoto similarity with sorafenib are provided. Less negative docking scores are preferred for ABCG2 binding (e.g., −7.0 is better than −8.0), indicating weaker binding. A Tanimoto similarity value between 0.4 and 0.7 suggests the ligand is novel yet exhibits structural similarity to sorafenib. This similarity suggests the potential for comparable biological activity. A lower molecular weight (ideally below 500 Da) is preferred, as it is associated with better drug-like properties. Low molecular weight ligands also tend to have better cell membrane permeability, improving their ability to reach the target site. Finally, we prioritize compounds with minimal lead-likeness violations to ensure favourable characteristics for drug development.

**Fig. 3.**
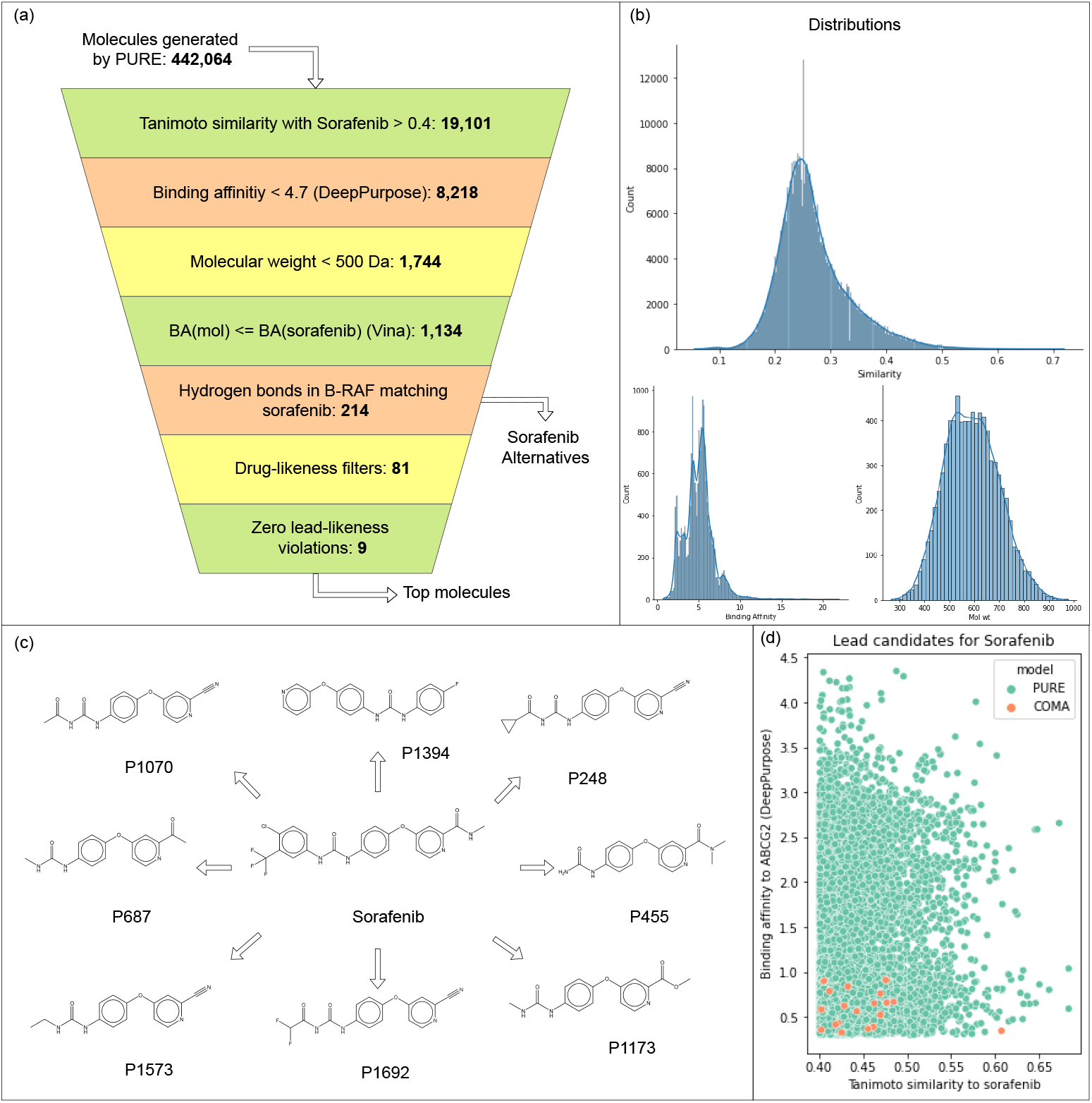
Filtration process to select lead candidates; (a) shows a funnel diagram for the number of molecules filtered at each step. (b) shows the distributions of molecules for the first three filtration steps. (c) shows 8 of the “top molecules” (d) shows a comparison of the 8,218 molecules that pass the first two filters against COMA molecules (these are the only filters used in COMA).

The fact that the Tanimoto similarity scores ranged from 0.4 to 0.7 indicates that the generated molecules are not only derived from the SCMG approach but also display novel chemical features. Lead-likeness analysis reveals that, while sorafenib itself does not meet all drug-likeness and lead-likeness parameters, many PURE molecules perform well. In conclusion, it can be inferred that PURE-generated molecules bind well to the desired target while also possessing ideal characteristics for further development in drug discovery based on sorafenib. Overall, this case study demonstrates that PURE can generate small molecules with high binding affinity, specific interactions, and favourable drug-like properties, making it a robust and reliable tool for SCMG compared to other algorithms like COMA.

## 3 Discussion

Deep learning has emerged as a powerful tool in drug discovery, but many existing methods struggle to address the complexities of the field effectively. Current SCMG approaches often introduce biases into their models, limiting the effectiveness of generated molecules *across* various evaluation metrics. Consequently, these molecules excel primarily in the optimization metrics but exhibit poor performance in practical applications as a result of metric bias and reduced generation diversity due to metric overfitting.

To combat metric bias, PURE is trained to learn an auxiliary path-prediction task under a similarity learning framework, to achieve structure-constrained molecular generation without relying on task-specific metrics. The framework inherently generates molecules with structural similarity to target molecules via a goal-conditional RL setting, which learns high-quality representations of state-action pairs. Under this framework, representations of similar state–action pairs are learned such that they are near each other in the representation space and their inner product translates to the RL value function [35]. This value function further captures the similarity between the state-action pairs [40, 41]. Thus, the inner product between the representations for two different state-action pairs gives a similarity score! This similarity score further pertains to the task being learned, therefore, in PURE, the similarity between state-action pairs is indicative that they lead to similar targets. This is one of the most important aspects of PURE, as it allows us to use the inner product of state-action representations as a surrogate for structural similarity.

Consequently, state-action pairs with the same state and different actions give similarity over actions. Therefore, actions that are applicable to a state and have similar representations lead to similar targets. This observation lets us sample top-*k* actions *closest* to the actor’s prediction, leading to an efficient generation process. During generation, PURE uses beam search to select the actions with the closest representations to the actor’s prediction to apply to the current molecule. It repeats for *n* steps to generate a large number of possible candidates.

Once the generation procedure is finished, a subset of molecules is selected that satisfies the desired combination of properties to be our final set of molecules. This way, the property metrics do not affect the training or generation procedure, and we obtain a single model under the PURE framework that can be evaluated on all property benchmarks. Thus, task-specific metrics are decoupled from model training.

As a result of decoupling the task-specific metrics from the training process, we reduce metric bias and increase the diversity of generated molecules. For all property benchmarks, PURE produces a diverse and competitive array of drug possibilities with high property and similarity scores. When optimizing for either property or similarity metrics, PURE obtains the best scores for the chosen metric in most benchmarks. When balancing both similarity and property, PURE gives competitive performance on all fronts, obtaining the best overall combined score across benchmarks. Notably, PURE offers the greatest diversity in the proposed molecules and significantly outperforms all other models in diversity and novelty across benchmarks. This result is particularly prominent since the average similarity amongst the generated molecules by the other methods is comparable to or higher than their similarity to the target molecule. This indicates that these models restrict themselves to a narrow search space and tend to generate molecules from that space. On the other hand, PURE explores a significantly larger space while achieving similar or higher performance on all metrics.

Furthermore, since our framework operates on molecular graphs using reaction rules, any action performed by the agent during the generation phase results in a valid molecule. Additionally, the model consistently produces novel and diverse molecules as a result of the auxiliary training of the agent. By starting the agent off with a variety of basic molecules, PURE can produce new molecules with similar structures that are fundamentally different from each other, leading to consistent novelty and diversity among outputs. The novelty observed in our results is due to our unique strategy of training the generation model on a dataset unrelated to the lead optimization dataset. By confining PURE to search within a discrete space through the iterative application of reaction rules, we not only encourage exploration that aligns with the natural process of chemical synthesis but also eliminate the complexity of generating valid and synthesizable molecules from the learning agent. In contrast, most generative methods for SCMG, which are fine-tuned using RL, are restricted to exploring only a small region around the base policy.

A drawback of PURE is its dependence on the availability of property metrics in the generation space. Since PURE is not trained to optimize these metrics, it does not inherently produce molecules with high scores in these areas. Instead, it generates many diverse molecules similar to the target and then selects those with the desired combinations. As seen in the DRD2 benchmark, PURE can struggle when the property for optimization is sparse. However, this trade-off allows for the flexibility of selecting the desired combination of similarity and property post-generation. For this reason, we argue that PURE is more useful for practical applications of SCMG than state-of-the-art methods, as the primary requirement in these cases is to optimize binding affinity. Finding a sizable dataset for training generative models for binding affinity optimization tasks is often challenging. Additionally, fine-tuning with RL for binding affinity is restrictive due to its slow computation.

To substantiate this claim, we further analyze PURE’s ability to generate sorafenib alternatives via the case study in section 2.2 compared to COMA. Despite fine-tuning for predicted binding affinity in COMA, or perhaps as a result of it, COMA generates a very small number of unique molecules that are further examined - 19 unique molecules out of 10,000 generated. On the other hand, PURE generates 24,290 molecules with Tanimoto similarity greater than 0.4 with sorafenib, out of which 19,101 molecules were unique. If the desired properties exist in this set of generated molecules, it is trivial to identify and filter such molecules. Here, we rely on the assumption that molecules similar to one another are also likely to exhibit some similar properties. Generating a large number of similar molecules would then result in molecules exhibiting a variety of similar properties, where a few would be amongst our targeted ones. The final goal of PURE then becomes to ensure that the set of generated molecules is similar to the target but is also diverse so that the desired properties can be found.

As a result of the rigorous filtering process and docking simulations, we are able to successfully identify a set of 214 molecules that closely resemble sorafenib in terms of their binding affinity to BRAF, molecular orientation within the active site, and interactions with key residues. These molecules, exhibiting a Tanimoto similarity greater than 0.4 with sorafenib and forming hydrogen bonds with all four critical residues in the BRAF active site, demonstrate promising potential as alternatives to sorafenib. With further analysis using stringent ADME filters, we can prioritize molecules with desirable properties such as drug-likeness, lead-likeness, and the absence of problematic structural features. Molecule descriptors, Tanimoto similarity value, docking scores, target-ligand interactions, and ADME properties of the 214 PURE ligands are present in Supplementary Table 1. A key finding is the identification of 30 PURE ligands with minimal violations in leadlikeness criteria. This is a significant improvement over the reference drug, sorafenib, which exhibits multiple violations. The top 9 molecules, with no violations, represent the most promising candidates for further development. Synthesis Pathways generated by PURE leading to these 9 molecules are present in Supplementary Figure 1.

Of course, to further validate these findings, it would be beneficial to conduct experimental studies to assess the biological activity and safety profiles of the identified molecules.

### 3.1 Conclusion

We have presented PURE, a novel framework for SCMG that departs from traditional approaches that are typically biased towards one or more metrics. By leveraging a policy-guided, metric-agnostic strategy, PURE enables the generation of drug-like molecules with high novelty, diversity, and synthesizability—all critical factors in real-world drug discovery. Our case study on sorafenib alternatives illustrates PURE’s potential to generate viable candidates capable of overcoming known drug resistance pathways, showcasing its practical utility in addressing the demanding challenges of modern drug discovery. Beyond this application, PURE’s RL-driven approach opens new possibilities in exploring expansive, discrete molecular spaces, paving the way for more innovative molecular design. As the field of AI-driven drug discovery evolves, PURE stands as a promising foundation for future advancements, with the potential to accelerate the discovery of therapeutic compounds across various domains. We envision PURE contributing not only to the identification of new drug-like molecules but also to the broader shift toward more adaptable, unbiased molecular design frameworks that can meet the demands of precision medicine and beyond.

## 4 Methods

In this section, we describe the design of PURE and its various components.

### 4.1 Problem setup

We observe that while SCMG may be modeled under the goal-conditioned reinforcement learning (GCRL) paradigm with similarity as a distance-based reward, its binary-reward equivalent would be to generate the target molecule itself (given some source molecule), i.e. source-to-target path prediction. Therefore, under a similarity learning framework, relaxation of source-to-target path prediction would give us SCMG. To realize this idea, we use the results from Eysenbach et al. [35] to design our method such that similar inputs occupy nearby regions in the embedding space and the inner product between their embeddings represents the similarity between the input pair. Therefore, input pairs that lead to the same target have similar representations, and hence, the model trained for source-to-target path prediction can be zero-shot transferred for SCMG while also being able to use the inner product between the representations as a surrogate for similarity during the generation procedure.

We first create an environment for molecular transitions using reaction templates and set up a source-to-target path prediction learning setup. Further, we make use of offline RL to overcome the sparse reward problem in binary-reward GCRL. Details about the RL environment and offline RL data generation procedure are described in Sections 4.2 and 4.4.

### 4.2 RL environment

#### State

The state is a tuple of two molecules: the current molecule and the target molecule (or goal) ⟨*s_t_, g*⟩.

#### Action

An action is a tuple of the reactant’s signature *r*^*sig*^ and the product’s signature *p*^*sig*^. *r*^*sig*^ and *p*^*sig*^ correspond to the functional groups removed and added during a chemical reaction, along with neighbors up to two atoms away, to denote the structures on which the rule is applicable. To determine the site of the reaction, the mining process also includes information about reaction centers, which are the atoms in the signatures where the subgraph addition or removal occurs. An example is shown in Figure 4. For this study, 84,968 reaction rules were mined from the USPTO-MIT dataset [36] according to the procedure described in Section 4.3 and used as the action set for our RL environment.

**Fig. 4.**
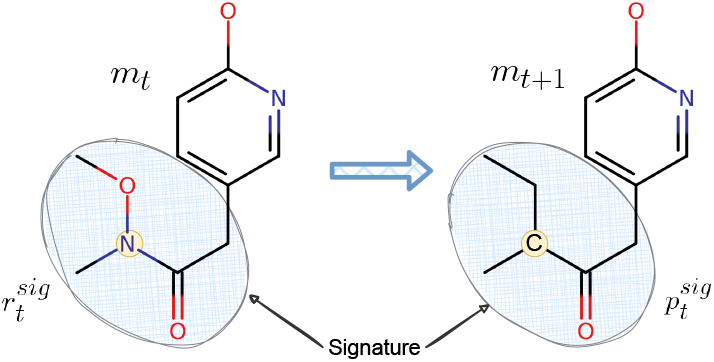
Reaction signatures: The illustration depicts reaction signatures involved in the reaction transformation of 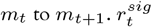 is the reactant signature and 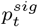 is the product signature. Centres are marked in yellow. The signatures are calculated as the changed subgraphs + neighbors up to 2 atoms away.

#### Transition

A direct replacement of *r*^*sig*^ on the reactant molecule with *p*^*sig*^ around the reaction centers is used to determine the product. Since the applicability of an action is determined by the reactant’s signature, given a reactant molecule, all those actions that have their reactant signature present in the molecule are considered to be applicable on it.

#### Rewards

We use binary rewards in our GCRL setting with a small penalty to negative samples to prevent zero PG loss when the return is zero. The reward function ℛ is defined as:

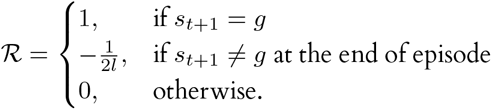

where *l* is the number of negative samples per positive sample, as described in section 4.4.2. Our choice of ℛ when *s*_*t* + 1_ ≠ *g* at the end of episode is experimental. We found that under the condition *G*(*a_t_* leads to *g*) ≤ *l* × *G*(otherwise), the actor’s performance starts to deteriorate. On investigation, we found that this is due to the cumulative magnitude of gradients from the negative return trajectories per positive trajectory being greater than that of the positive return trajectory.

#### State and action representations

States (molecules) and actions (signatures) are graph objects. Hence, we use a Graph Isomorphism Network (GIN) [42] to encode each and create learnable representations.

### 4.3 Mining Reaction Rules

We adapt the methodology described in [43] for mining reaction rules. We represent molecules as graphs, with atoms as vertices and bonds as edges. We then establish a reactant-product mapping for transformed metabolites based on their closest molecular weights. Through a scalable graph mining procedure[44], we identify critical subgraphs for reactions, termed reaction signatures. For instance, during the oxidation of alcohols by alcohol dehydrogenase, the *CH*_2_*OH* subgraph serves as the reaction signature, with the corresponding reaction rule as *CH*_2_*OH* converting to *CHO*.

We adapt the above procedure to extract reaction rules from the USPTO-MIT dataset [36]. We first establish a one-to-one mapping between reactants and products based on molecular weight proximity. This helps to identify the most likely transformation pairs [43]. For instance, in the reaction: *Acetyl chloride + Ammonia* →*Acetamide + Hydrogen chloride*, Acetyl chloride would be mapped to Acetamide. From this extracted set of reactant–product pairs, duplicates are removed to ensure that only unique transformations are considered. Subsequently, reaction centers and signatures are extracted by subtracting the maximum common substructure from each reactant–product pair. This step helps to identify the specific regions of the molecules that undergo significant chemical changes during the reaction. An example of reaction centers and signatures is shown in Figure 4. Finally, we retain the reactions with a single reaction center, as these provide more concise and reliable rules. While reactions with multiple centers do exist, they often represent complex or ambiguous transformations.

### 4.4 Offline RL data

#### 4.4.1 Positive samples

We generate an offline RL dataset using trajectories with high rewards to overcome the sparse reward problem in GCRL. However, given an arbitrary source–target pair, finding if it has a high reward trajectory requires us to know whether there exists a path between the two molecules, which is non-trivial. Instead, if some policy is used to roll out a trajectory, by virtue of the creation of the trajectory, there exists a path from its starting molecule to the final molecule. The trajectory would, therefore, have a high reward, making it a valid trajectory for the dataset. Following this idea, we select a starting molecule and use a uniform random policy to perform rollouts over *n* steps to create an *n*-step offline RL dataset.

We start by selecting a source molecule and performing a rollout for *n* steps using a uniform random policy. This generates a trajectory ⟨*m*_0_, *rr*_0_, *m*_1_, *rr*_1_,…, *m*_*n−*1_, *rr*_*n*−1_, *m_n_*⟩, where *m_t_* represents the molecule at step *t* and *rr_t_* is the reaction rule applied on *m_t_* to create *m*_*t*+1_. Since every *m_t_* has a path to *m_n_* (*t* ≠ *n*), every sub-trajectory ⟨ *m_t_, rr_t_, m*_*t*+1_, *rr*_*t*+1_,…, *m*_*n*−1_, *rr*_*n−*1_, *m_n_*⟩, is also a valid trajectory.

Therefore, for each *t* ∈ [0, *n* −1], we save ⟨*s_t_, a_t_, s*_*t*+1_, *r*⟩ as the state, action, next state and reward; where the state is a tuple containing the molecule at that state and the target molecule i.e. *s_t_* = (*m_t_, m_n_*), action *a_t_* = *rr_t_* and the reward *r* = *R*(*s_t_, a_t_, s_t_*+1) is calculated as described in the section 4.2. This constructs our offline RL dataset that contains positive or high-return samples.

#### 4.4.2 Negative samples

The offline dataset only contains trajectories with high returns. Hence for each sample in the training batch, we uniformly randomly sample *l* actions during runtime to constitute *negative samples*. These *l* actions are selected from the list of all actions except the action with a high return. Dynamically allowing the agent to see samples with low returns enhances the agent’s distinguishing capabilities and improves performance.

### 4.5 Training data

We generate an offline RL dataset with trajectories of length 5 as described above, containing 100,000 samples (20,000 source-target pairs), which we further subdivide into 80,000 and 20,000 for training and validation.

### 4.6 Loss

Due to our small episode length, we consider the un-discounted case. Using the reward function described in section 4.2, we can compute the return G as follows:

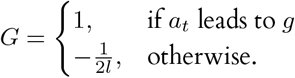

We use the standard policy gradient (PG) loss for actor *π_θ_* and MSE for the critic *Q_ϕ_*:

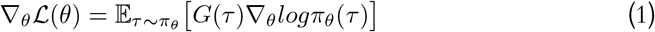

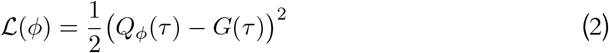

Since our environment is deterministic, we use the return in the PG loss instead of the critic’s estimate. We still train a critic network, as it is responsible for learning the policy-guided representations [35]. Moreover, we use the critic’s estimate in the beam search during generation.

### 4.7 Algorithms

#### 4.7.1 Training

Algorithm 1 describes the pseudocode for the training algorithm. Figures 5(a), (b), and (c) show pictorial descriptions of the training algorithm. It is broadly divided into the following three steps:

**Fig. 5.**
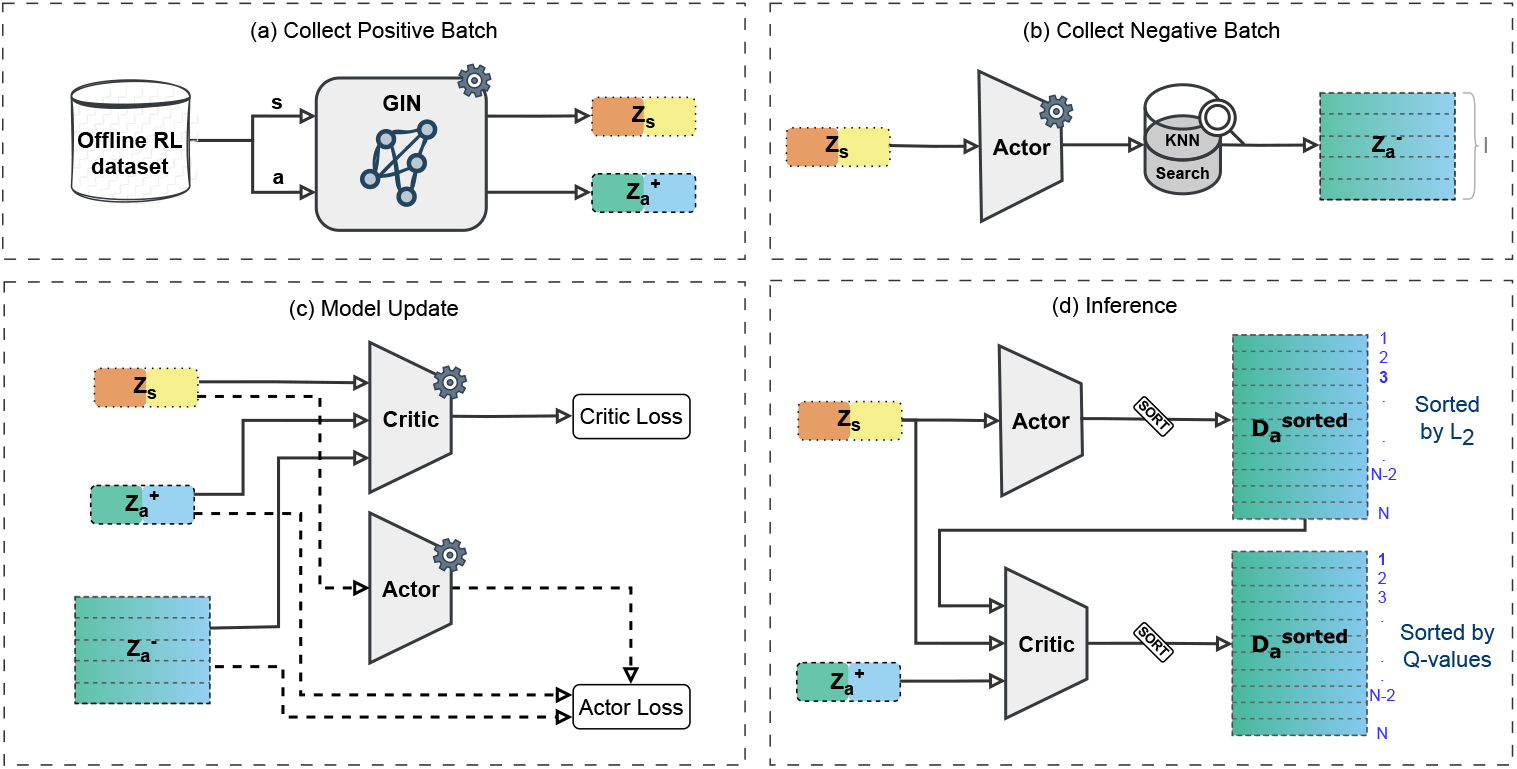
Training and evaluation procedure for actor-critic: (a) Sampling of positive samples from the offline RL dataset. The state *s* and corresponding correct action *a* are passed through the embedding module to create *Z_s_* and 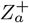, the state and positive action representations. (b) Creation of negative samples by randomly sampling action embeddings other than 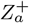. (c) Inputs to the actor and critic losses — the actor loss is calculated over the action embeddings 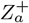 and 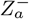, and the critic loss is calculated over the Q value. The gradients are then used to update the two models. (d) The evaluation procedure. First, the actor’s prediction is used to order the action dataset according to the Euclidean distance from the prediction to create to select the top *B_A_* actions. These are sent for re-evaluation to the critic. These are sorted by the critic’s Q values, and the top *B* actions are returned.

1. *Collect positive batch*: In this step, a batch of transitions is sampled from the offline RL dataset.
2. *Collect negative batch*: A negative batch is sampled using the strategy described in section 4.4.2.
3. *Model update*: The positive and negative batches are passed through the actor and critic models to calculate their respective losses and backpropagated to update their parameters.

##### Algorithm 1

Pseudocode for training actor-critic

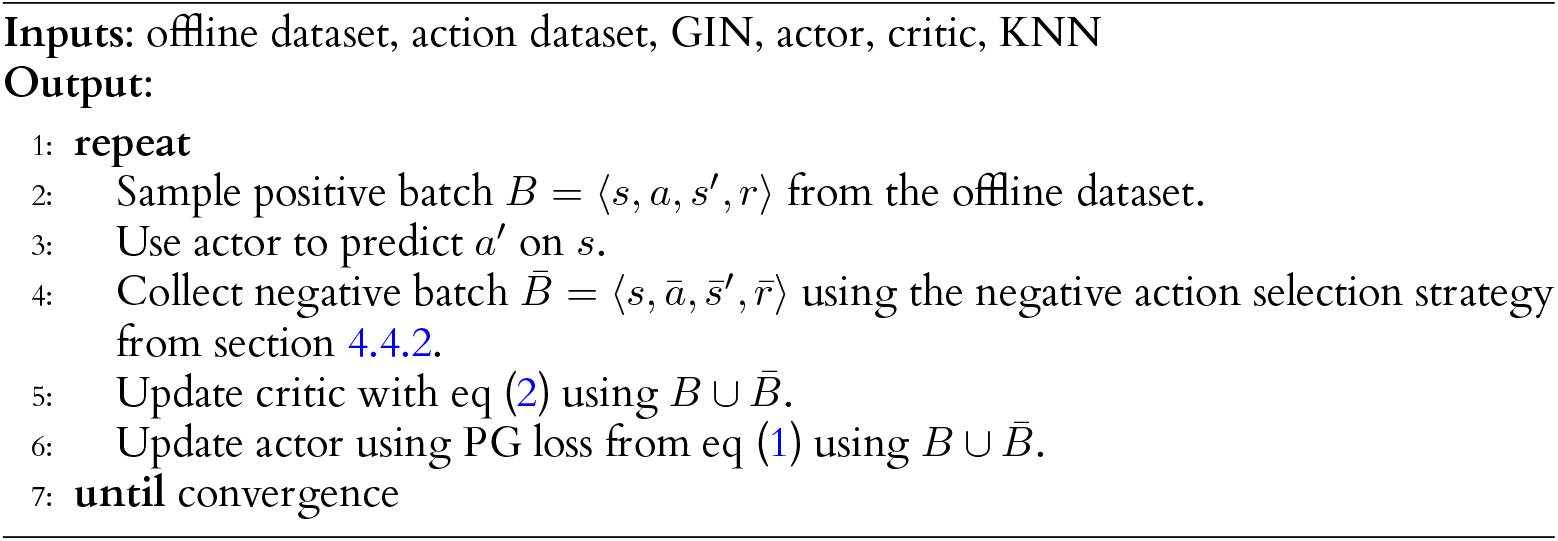

##### Algorithm 2

Pseudocode for generating molecules similar to target molecule

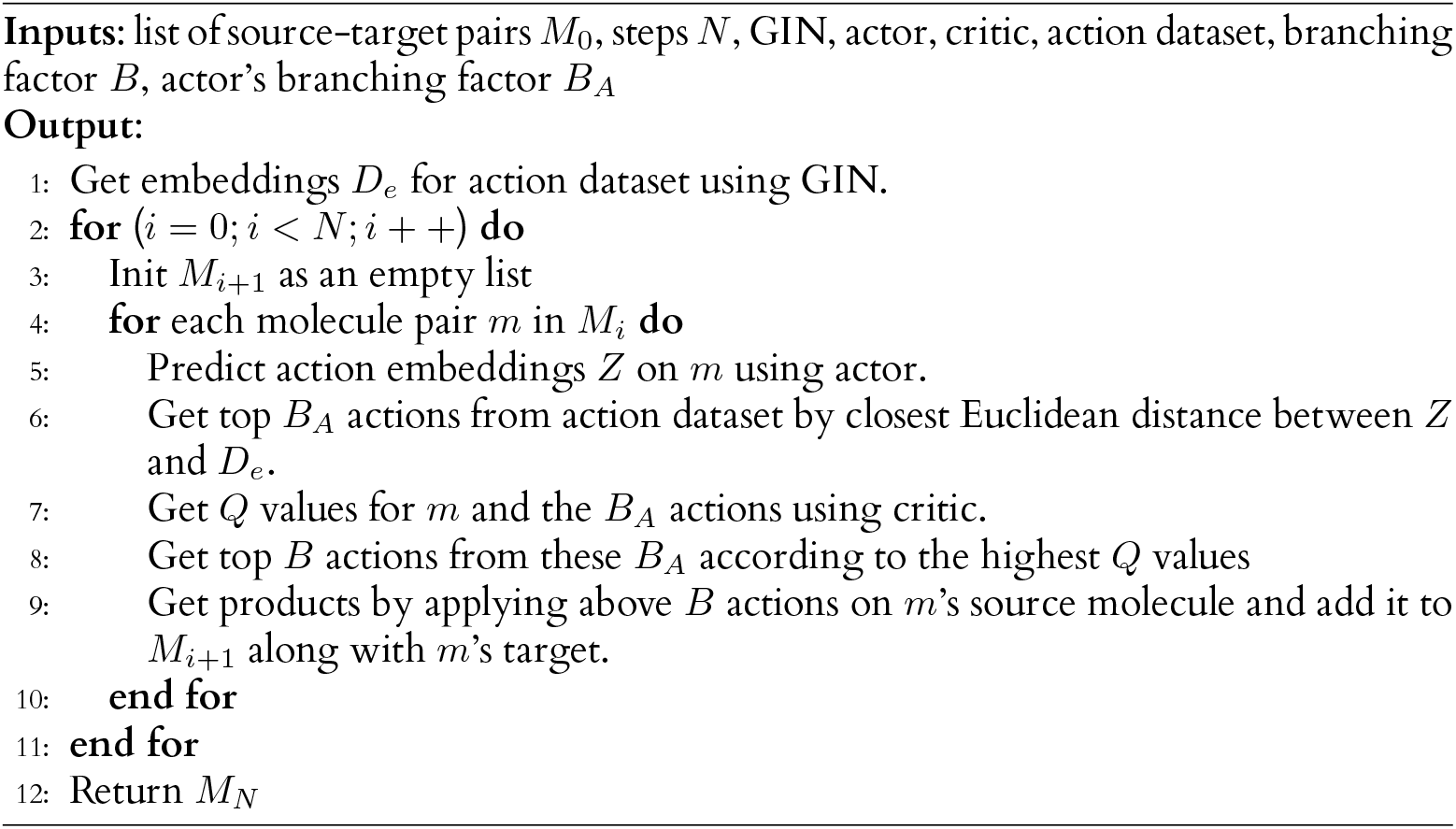

#### 4.7.2 Generation

Algorithm 2 describes the pseudocode for generating molecules similar to the target molecule. Figure 5(d) shows the generation procedure. It is broadly divided into the following three steps:

1. *Filter actions using actor*: Get the closest actions according to Euclidean distance from the actor’s prediction.
2. *Filter actions using critic*: Get the actions with the highest *Q* values according to the critic.
3. *Apply*: Apply the filtered actions on the current source to get the next list of sources (or final candidates if in the last iteration).

### 4.8 Evaluation

During evaluation, test data contains only target molecules. Starting molecules are selected from the Enamine Building Block catalogue^1^ Global Stock. They are filtered using the highest similarity between the starting molecules and targets, the highest similarity between the scaffolds (using rdkit’s *GetScaffoldForMol* function) of starting molecules and targets, and the maximum common subsequence (using rdkit’s *FindMCS* function) between starting molecules and targets to select 30 unique starting molecules per target. The generation procedure is then run on all 30 pairs as described in Section 4.7 for *N* = 5 time steps, branching factor *B* = 5, and actor’s branching factor *B_A_* = 50. Following Choi et al. [3], we select 20 molecules per target for our evaluation. Due to the nature of our generation process, we sample subsets from our generated molecules for the desired properties. We select molecules with the highest (*x similarity* + *y property*) and choose different values of *x* and *y* to sample different subsets.

Notably, the generation process is not repeated for sampling different subsets. The generation process is run once for each target, and the subsets are sampled post-generation. Additionally, we do not train different models for different criteria. The same model is used to evaluate *all* the benchmarks and coefficient combinations.

#### 4.8.1 Benchmarks

We evaluate PURE on four different benchmark datasets - QED, DRD2, pLogP04 and pLogP06 from [45]. The test benchmark for QED contains 800 molecules with the task to generate molecules similar to those in the test data but with higher QED. The test benchmark for DRD2 contains 1,000 molecules with the aim to generate similar molecules with improved biological activity against the dopamine type 2 receptor (DRD2). The test benchmarks for pLogP04 and pLogP06 contain 800 molecules each with the aim tointending to generate similar molecules with better plogP^2^.

We evaluate PURE on the above benchmarks and show comparisons against the results in tables S3, S5, S7 and S9^3^ in Choi et al. [3].

#### 4.8.2 Baselines

The following methods are used as baselines to evaluate the performance of PURE: [3], JTVAE [11], VJTNN, VJTNN+GAN [12], CORE [13], HierG2G [14], HierG2G+BT [15], and UGMMT [16].

- COMA learns structural similarity by employing contractive and margin losses and fine-tuning for property optimization using RL.
- JTVAE generates a tree representing the molecules’ substructure using an encoder– decoder architecture and then uses a message passing network to combine the substructures into a molecule.
- VJTNN is an extension of JTVAE by adding an attention mechanism to the decoder.

VJTNN+GAN further integrates an adversarial component to the training.

- CORE uses a copy-and-refine strategy to decide whether to copy a substructure from the input or to generate a new substructure.
- HierG2G generates molecular graphs using structural motifs as building blocks with a hierarchical encoder-decoder architecture.
- HierG2G+BT extends HierG2G with additional synthetic training data generated using a semi-supervised technique called back translation.
- UGMMT uses a double-cycle training scheme for molecular translation to generate molecular SMILES.

##### Metrics

We evaluate our method on the metrics from COMA [3], as given below:

1. Total score: The mean of the below six metrics
2. Validity: The ratio of generated molecules that are valid
3. Property: The average of (desired) property scores of the generated molecules
4. Improvement: The average of the difference of property scores between valid generated molecules and source molecules of test data
5. Similarity: The average of the Tanimoto similarity between valid generated molecules and source molecules of test data
6. Novelty: The ratio of valid generated molecules that are not in the training data
7. Diversity: The average of Tanimoto dissimilarity (= 1 - Tanimoto similarity) between generated molecules

### 4.9 Filtering Molecules in the Sorafenib Case Study

Initially, the entire set of unique molecules generated by PURE is sequentially filtered using two parameters: their similarity to sorafenib and their binding affinity towards ABCG2, as computed by DeepPurpose [46]. Subsequently, the pool is further refined to identify ligands with a molecular weight less than 500 Da, consistent with Lipinski’s Rule of 5, a molecular weight cutoff suitable for identifying lead-like and orally bioavailable small molecules. Figure 3(a) shows the filtration steps from start to end along with the number of molecules obtained at each step. A total of 442,064 unique molecules are generated by PURE, out of which 19,101 molecules exhibit a Tanimoto similarity greater than 0.4 with sorafenib. From this set, 8,218 molecules are selected that exhibited a binding affinity towards ABCG2 lesser than 4.7. Finally, 1,744 molecules of molecular weight under 500 Da are chosen to be carried forward for docking simulations.

### 4.10 Docking Simulations for the Sorafenib Case Study

Docking simulations against both BRAF and ABCG2 are carried out for each of the 1,744 initially filtered sorafenib alternatives generated by PURE. The 3-D structures of all ligands are prepared from their respective SMILES notations using RDKit v. 2024.03.1 [47] and Open Babel v. 3.1.0 [48]. The prepared molecules are energy-minimised and generated in PDBQT format for docking using Autodock Vina v. 1.2.5 (hereafter referred to as Vina) [49]. The X-ray diffraction structure of BRAF (PDB ID: 1UWH) and cryo-EM structure of ABCG2 (PDB ID: 6VXH) are obtained from RCSB PDB and used as the receptors for the docking simulations. The receptor structures are prepared for docking using MGLTools v. 1.5.7 [50], and the docking grid parameters are generated using the active-site residues.

Molecular docking simulations are carried out for all ligands against both receptors at an exhaustiveness parameter of 40. 20 binding poses are generated for each docked ligand. The top binding pose with the best docking score of each molecule is carried forward for further analysis. The complexes are visualised using PyMOL v. 3.0.2 [51]. Following this, the tool PLIP v. 2.3.0 (Protein Ligand Interaction Profiler) [52] is used to generate the interaction profiles of the protein-ligand complexes. PLIP identifies various types of interactions between proteins and ligands, including hydrogen bonds, hydrophobic interactions, *π*−*π* stacking, cation-*π* interactions, halogen bonds and salt bridges. Analysing these interactions provides further insights into the binding affinity and specificity of ligands. The docking scores obtained from Vina, the PLIP results and the binding poses of the ligands within the active site are used to evaluate the lead-like qualities of the PURE and COMA ligands, using sorafenib as a reference. Ligands that exhibit properties closer to that of sorafenib are considered promising candidates and inspected further in detail.

Specifically, the docked ligands are evaluated based on two criteria: their binding affinity, represented by the Vina docking score towards ABCG2, which has to be equal to or lower than that of sorafenib’s affinity towards ABCG2, and their ability to replicate a hydrogen bond network formed by sorafenib with BRAF. Molecules that form fewer hydrogen bonds are rejected, even if they exhibit other interactions. However, those that show additional interactions beyond these specific hydrogen bonds are retained.

The final step in the process is an ADME analysis of the molecules from the H-bond filtration step using the SwissADME server [53]. The SMILES of sorafenib and molecules that pass the previous step are used as input to SwissADME and are evaluated based on various drug-likeness and pharmacokinetic properties to ensure that the top molecules to be selected adhere to medicinal chemistry standards expected of lead-like molecules. Furthermore, the Tanimoto similarities of the passing molecules with sorafenib are also noted.

## Declarations

### Availability of Data and Materials

Our data and code is publicly available at https://github.com/MrScabbyCreature/ReactionRL.

### Authors’ Contributions

A.G., B.R., K.R., and S.P. conceived the original study. A.G., S.C., B.R., R.B., K.R., and S.P. co-designed the methodology, which was subsequently refined and implemented by A.G. and experimentally tested and evaluated by A.G. and S.C. B.L. and A.G. led the evaluation of the case study, assisted by the other authors. All authors jointly contributed to the data analysis and writing of the manuscript led by A.G., K. R., and S.P.

## Acknowledgments

Not applicable.

## Footnotes

1 https://enamine.net/building-blocks

2 The pLogP04 and pLogP06 benchmarks differ in the nature of the creation of their training dataset but we do not use the training data for these benchmarks.

3 Due to the nature of our generation process, a comparison of success rates against tables S2, S4, S6 and S8 is unsuited.

