## Supplementary Table 1 for "PURE: Policy-guided Unbiased REpresentations for structure-constrained molecular generation"

Supplementary Table 1: Molecule descriptors, Tanimoto similarity value, docking scores, target-ligand interactions, and ADME properties of 214 PURE ligands

| S. No. | Ligand | Canonical SMILES | Molecular weight (Da) | Tanimoto Similarity | Docking score with BRAF (kcal/mol) | Docking score with ABCG2 (kcal/mol) | Hydrophobic interactions | Hydrogen bonds | Salt bridge | Pi stack interaction(s) | Pi-cation interaction | Hydrogen bond match | Lipinski violation(s) | Ghose violation(s) | Veber violation(s) | Egan violation(s) | Muegge violation(s) | Bioavailability score | PAINS alert(s) | Brenk alert(s) | Leadlikeness violation(s) | Synthetic accessibility score |
|---|---|---|---|---|---|---|---|---|---|---|---|---|---|---|---|---|---|---|---|---|---|---|
| 1 | P14 | CNc1ccc(cc1C(F)(F)F)NC(=O)Nc1ccc(cc1)Oc1cncc(c1)Cl | 436.81 | 0.65 | -10.974 | -8.117 | GLU 500, LEU 504, LEU 513, THR 528, TRP 530, ASP 593 | LYS 482, GLU 500, CYS 531, ASP 593 | - | PHE 594 | - | Yes | 0 | 1 | 0 | 1 | 0 | 0.55 | 0 | 0 | 3 | 3.02 |
| 2 | P15 | O=C(Nc1ccc(c(c1)C(F)(F)F)Cl)Nc1ccc(cc1)Oc1cnccc1C | 421.8 | 0.68 | -11.679 | -8.401 | VAL 470, LYS 482, LEU 504, LEU 513, THR 528, ASP 593 | LYS 482, GLU 500, CYS 531, ASP 593 | - | PHE 594 | HIS 573 | Yes | 0 | 1 | 0 | 1 | 0 | 0.55 | 0 | 0 | 2 | 2.83 |
| 3 | P17 | O=C(Nc1ccc(c(c1)C(F)(F)F)N)Nc1ccc(cc1)Oc1ccncc1 | 388.34 | 0.62 | -11.16 | -8.177 | ALA 480, LYS 482, VAL 503, LEU 504, LEU 513, THR 528 | LYS 482, GLU 500, CYS 531, ASP 593 | - | PHE 594 | HIS 573 | Yes | 0 | 1 | 0 | 1 | 0 | 0.55 | 0 | 1 | 1 | 2.52 |
| 4 | P20 | O=C(Nc1ccc(c(c1)C(F)(F)F)Cl)Nc1ccc(cc1)Oc1cncc(c1)Cl | 442.22 | 0.68 | -11.661 | -8.246 | ALA 480, LYS 482, VAL 503, LEU 504, LEU 513, THR 528 | LYS 482, GLU 500, CYS 531, ASP 593 | - | PHE 594 | HIS 573 | Yes | 0 | 1 | 0 | 1 | 1 | 0.55 | 0 | 0 | 2 | 2.81 |
| 5 | P22 | COC(=O)c1ccc(nc1)c1cc(ccc1N(C)C)NC(=O)Nc1ccc(cc1)Oc1cnccc1C | 497.55 | 0.41 | -10.857 | -8.245 | ILE 462, VAL 470, ALA 480, GLU 500, VAL 503, LEU 504, THR 528, ASP 593, PHE 594 | LYS 482, GLU 500, CYS 531, ASP 593 | ARG 574 | TRP 530, PHE 594 | - | Yes | 0 | 2 | 0 | 0 | 0 | 0.55 | 1 | 0 | 3 | 3.72 |
| 6 | P31 | O=C(c1nccc(c1)Oc1ccc(cc1)N(O)C)NCc1cccc(c1)C(=O)O | 393.39 | 0.44 | -9.452 | -8.409 | ALA 480, LYS 482, THR 528 | LYS 482, GLU 500, CYS 531, SER 535, ASN 579, ASP 593 | - | PHE 582, PHE 594 | - | Yes | 0 | 0 | 0 | 0 | 0 | 0.56 | 0 | 1 | 2 | 2.77 |
| 7 | P34 | COC(=O)c1nccc(c1)Oc1ccc(cc1)NC(=O)NCCOc1ccc(cc1N(C)C)[N+](=O)[O-] | 495.48 | 0.45 | -9.821 | -7.945 | LYS 482, VAL 503, THR 528 | LYS 482, GLU 500, CYS 531, ASP 593 | - | PHE 594 | - | Yes | 1 | 2 | 2 | 1 | 0 | 0.55 | 0 | 2 | 2 | 3.73 |
| 8 | P44 | O=C(Nc1ccc(cc1)Oc1ccnc(c1)C(=O)CC(=O)C)NC(=O)C1CC1 | 381.38 | 0.43 | -10.153 | -7.883 | ILE 462, LYS 482, GLU 500, VAL 503, LEU 504, LEU 513, THR 528, PHE 582, PHE 594 | LYS 482, GLU 500, CYS 531, ASP 593 | - | PHE 594 | - | Yes | 0 | 0 | 0 | 0 | 0 | 0.55 | 0 | 1 | 2 | 2.73 |
| 9 | P46 | ClCc1ccc(cc1)NC(=O)Nc1ccc(cc1)Oc1cncc(c1)S | 385.87 | 0.45 | -10.047 | -7.439 | LYS 482, GLU 500, VAL 503, LEU 504, LEU 513, THR 528 | LYS 482, GLU 500, CYS 531, ASP 593 | - | PHE 594 | - | Yes | 0 | 0 | 0 | 0 | 0 | 0.55 | 0 | 2 | 1 | 2.78 |
| 10 | P56 | COC(=O)NC(=O)Nc1ccc(cc1)Oc1ccnc(c1)CN | 316.31 | 0.44 | -8.295 | -7.683 | ALA 480, LYS 482, TRP 530, PHE 594 | LYS 482, GLU 500, CYS 531, ASP 593 | - | PHE 594 | - | Yes | 0 | 0 | 0 | 0 | 0 | 0.55 | 0 | 0 | 1 | 2.42 |
| 11 | P72 | NCc1ccc(cc1)NC(=O)CNC(=O)Nc1ccc(cc1)Oc1ccnc(c1)C#N | 416.43 | 0.47 | -10.029 | -8.158 | ALA 480, LYS 482, VAL 503, THR 528, ILE 571 | LYS 482, GLU 500, CYS 531, ILE 572, ASP 593 | - | PHE 594 | - | Yes | 0 | 0 | 1 | 1 | 0 | 0.55 | 0 | 0 | 2 | 3.01 |
| 12 | P73 | COC(=O)NC(=O)Nc1ccc(cc1)Oc1ccnc(c1)C(=O)N(C)C | 358.35 | 0.45 | -9.952 | -7.545 | ALA 480, LYS 482, LEU 513, THR 528 | LYS 482, GLU 500, CYS 531, ASP 593 | - | PHE 594 | - | Yes | 0 | 0 | 0 | 0 | 0 | 0.55 | 0 | 0 | 2 | 2.62 |
| 13 | P76 | CCc1ccncc1Oc1ccc(cc1)NC(=O)Nc1ccc(cc1)C(C(=S)N)(C)C | 434.55 | 0.41 | -10.316 | -8.163 | ILE 462, VAL 470, ALA 480, LYS 482, GLU 500, VAL 503, LEU 504, THR 528, LEU 566, ILE 571, ASP 593 | LYS 482, GLU 500, CYS 531, HIS 573, ASP 593 | - | TRP 530, PHE 594 | - | Yes | 0 | 0 | 0 | 0 | 0 | 0.55 | 0 | 1 | 3 | 3.22 |
| 14 | P81 | N#Cc1nccc(c1)Oc1ccc(cc1)NC(=O)NC(=O)Oc1ccccc1 | 374.35 | 0.44 | -10.394 | -8.045 | ALA 480, LYS 482, VAL 503, LEU 513, THR 528, ASP 593 | LYS 482, GLU 500, CYS 531, ASP 593 | - | HIS 573, PHE 594 | - | Yes | 0 | 0 | 0 | 0 | 0 | 0.55 | 0 | 0 | 3 | 2.9 |
| 15 | P85 | O=C(Nc1ccc(cc1)Oc1ccnc(c1)C(=O)C)NC(=O)OS(=O)(=O)c1ccccc1N | 470.46 | 0.46 | -11.189 | -8.189 | ILE 462, ALA 480, LYS 482, VAL 503, LEU 513, THR 528 | LYS 482, GLU 500, CYS 531, ASP 593 | HIS 573 | PHE 594 | - | Yes | 1 | 0 | 1 | 1 | 1 | 0.55 | 0 | 2 | 2 | 3.39 |
| 16 | P88 | NCCc1ccc(cc1)NC(=O)Nc1ccc(cc1)Oc1cncc(c1)C(=O)NC(C)C | 433.5 | 0.45 | -10.499 | -8.314 | ILE 462, LYS 482, GLU 500, VAL 503, LEU 504, LEU 513, THR 528, TRP 530, PHE 582 | LYS 482, GLU 500, CYS 531, ASP 593 | - | PHE 594 | - | Yes | 0 | 0 | 1 | 0 | 0 | 0.55 | 0 | 0 | 2 | 3.23 |
| 17 | P91 | C=COC(=O)NC(=O)Nc1ccc(cc1)Oc1ccnc(c1)C=O | 327.29 | 0.41 | -8.633 | -7.426 | ALA 480, GLU 500, VAL 503, LEU 504, THR 528, TRP 530 | LYS 482, GLU 500, CYS 531, ASP 593 | - | PHE 594 | - | Yes | 0 | 0 | 0 | 0 | 0 | 0.55 | 0 | 2 | 1 | 2.57 |
| 18 | P96 | O=C(Nc1ccc(cc1)Oc1ccnc(c1)C(=O)N(C)C)NC(=O)OCc1ccccc1 | 434.44 | 0.45 | -11.148 | -8.426 | ALA 480, LYS 482, VAL 503, LEU 513, LEU 566, ILE 571 | LYS 482, GLU 500, CYS 531, ASP 593 | - | TRP 530, PHE 594 | - | Yes | 0 | 0 | 1 | 0 | 0 | 0.55 | 0 | 0 | 2 | 3.15 |
| 19 | P103 | N#Cc1ccc(n1C)c1cc(ccc1N(C)C)NC(=O)Nc1ccc(cc1)Oc1cccnc1 | 452.51 | 0.41 | -10.307 | -7.818 | ALA 480, GLU 500, VAL 503, LEU 504, THR 528, ASP 593, PHE 594 | LYS 482, GLU 500, CYS 531, ASP 593 | - | TRP 530, PHE 594 | - | Yes | 0 | 1 | 0 | 0 | 0 | 0.55 | 1 | 0 | 3 | 3.64 |
| 20 | P104 | CCOC(=O)NC(=O)Nc1ccc(cc1)Oc1ccnc(c1)CN | 330.34 | 0.43 | -8.29 | -7.857 | LYS 482, GLU 500, LEU 513, THR 528 | LYS 482, GLU 500, CYS 531, ASP 593 | - | PHE 594 | - | Yes | 0 | 0 | 0 | 0 | 0 | 0.55 | 0 | 0 | 1 | 2.61 |
| 21 | P106 | O=CCCNC(=O)Nc1ccc(cc1)Oc1ccnc(c1)C(=O)N1CCCC1 | 382.41 | 0.4 | -9.432 | -7.278 | ILE 462, ALA 480, LYS 482, GLU 500, LEU 504, THR 528 | LYS 482, GLU 500, CYS 531, ASP 593 | - | PHE 594 | - | Yes | 0 | 0 | 0 | 0 | 0 | 0.55 | 0 | 1 | 2 | 2.74 |
| 22 | P131 | COC(=O)c1nccc(c1)Oc1ccc(cc1)NC(=O)NC(=O)c1ccccc1F | 409.37 | 0.51 | -10.875 | -8.082 | ALA 480, LYS 482, GLU 500, VAL 503, LEU 513, THR 528, ASP 593 | LYS 482, GLU 500, CYS 531, ASP 593 | - | PHE 594 | - | Yes | 0 | 0 | 0 | 0 | 0 | 0.55 | 0 | 0 | 3 | 2.86 |
| 23 | P134 | NCCc1ccc(cc1)NC(=O)Nc1ccc(cc1)Oc1cncc(c1)C(=O)N | 391.42 | 0.45 | -10.535 | -8.01 | LYS 482, GLU 500, VAL 503, LEU 504, LEU 513, THR 528, TRP 530, ILE 571 | LYS 482, GLU 500, CYS 531, HIS 573, ASP 593 | - | PHE 594 | - | Yes | 0 | 0 | 0 | 1 | 0 | 0.55 | 0 | 0 | 2 | 2.9 |
| 24 | P139 | ClCc1ccc(cc1)NC(=O)CNC(=O)Nc1ccc(cc1)Oc1ccnc(c1)C#N | 435.86 | 0.48 | -10.183 | -8.111 | ALA 480, LYS 482, VAL 503, THR 528, ILE 571 | LYS 482, GLU 500, CYS 531, ASP 593 | - | PHE 594 | - | Yes | 0 | 0 | 0 | 0 | 0 | 0.55 | 0 | 1 | 2 | 2.97 |
| 25 | P163 | CCOC(=O)NCc1cccc(c1)CNC(=O)c1nccc(c1)Oc1ccc(cc1)Nc1ccncc1 | 497.55 | 0.43 | -9.972 | -8.3 | ALA 480, LYS 482, PHE 497, GLU 500, LEU 513, THR 528, ASP 593, PHE 594 | LYS 482, GLU 500, CYS 531, ASP 593 | - | PHE 594 | - | Yes | 0 | 2 | 1 | 0 | 0 | 0.55 | 0 | 0 | 3 | 3.52 |
| 26 | P166 | N#Cc1nccc(c1)Oc1ccc(cc1)NC(=O)NC(=O)c1ccccc1Cl | 392.8 | 0.51 | -10.832 | -8.155 | ALA 480, LYS 482, GLU 500, VAL 503, LEU 504, LEU 513, THR 528 | LYS 482, GLU 500, CYS 531, ASP 593 | - | PHE 594 | - | Yes | 0 | 0 | 0 | 0 | 0 | 0.55 | 0 | 0 | 2 | 2.61 |
| 27 | P177 | COC(=O)c1nccc(c1)Oc1ccc(cc1)NC(=O)NC(=O)Cc1ccc(c(c1)OC)N | 450.44 | 0.48 | -10.206 | -8.121 | ALA 480, LEU 504, THR 528, ASP 593 | LYS 482, GLU 500, CYS 531, HIS 573, ASP 593 | - | TRP 530, PHE 594 | - | Yes | 0 | 0 | 2 | 1 | 0 | 0.55 | 0 | 1 | 2 | 3.2 |
| 28 | P179 | CNC(=O)CCc1ccncc1Oc1ccc(cc1)NC(=O)Nc1ccc(cc1)C1(C#N)CC1 | 455.51 | 0.42 | -10.605 | -7.981 | ILE 462, VAL 470, ALA 480, LYS 482, GLU 500, VAL 503, LEU 504, THR 507, THR 528, LEU 566, PHE 582, ASP 593 | LYS 482, GLU 500, CYS 531, ASP 593, PHE 594 | - | TRP 530, PHE 594 | - | Yes | 0 | 0 | 1 | 0 | 0 | 0.55 | 0 | 0 | 2 | 3.27 |

Supplementary Table 1: Molecule descriptors, Tanimoto similarity value, docking scores, target-ligand interactions, and ADME properties of 214 PURE ligands

| S. No. | Ligand | Canonical SMILES | Molecular weight (Da) | Tanimoto Similarity | Docking score with BRAF (kcal/mol) | Docking score with ABCG2 (kcal/mol) | Hydrophobic interactions | Hydrogen bonds | Salt bridge | Pi stack interaction(s) | Pi-cation interaction | Hydrogen bond match | Lipinski violation(s) | Ghose violation(s) | Veber violation(s) | Egan violation(s) | Muegge violation(s) | Bioavailability score | PAINS alert(s) | Brenk alert(s) | Leadlikeness violation(s) | Synthetic accessibility score |
|---|---|---|---|---|---|---|---|---|---|---|---|---|---|---|---|---|---|---|---|---|---|---|
| 29 | P180 | N#Cc1nccc(c1)Oc1ccc(cc1)NC(=O)NCC(=O)OC(C)(C)C | 368.39 | 0.41 | -9.289 | -6.972 | LYS 482, VAL 503, LEU 513, THR 528 | LYS 482, GLU 500, CYS 531, ASP 593 | HIS 573 | PHE 594 | - | Yes | 0 | 0 | 0 | 0 | 0 | 0.55 | 0 | 0 | 2 | 2.86 |
| 30 | P184 | ClCCCc1ccccc1CNC(=O)c1nccc(c1)Oc1ccc(cc1)N(C(=O)NC)O | 468.93 | 0.43 | -9.608 | -8.405 | ALA 480, LEU 513, THR 528, PHE 582, PHE 594 | LYS 482, GLU 500, CYS 531, ASP 593 | - | PHE 594 | - | Yes | 0 | 0 | 1 | 0 | 0 | 0.55 | 0 | 3 | 2 | 3.22 |
| 31 | P191 | CC(=O)CNC(=O)Nc1ccc(cc1)Oc1ccnc(c1)C(O)C | 329.35 | 0.43 | -9.252 | -7.698 | ILE 462, ALA 480, LYS 482, GLU 500, VAL 503, LEU 504, LEU 513, THR 528, TRP 530 | LYS 482, GLU 500, CYS 531, ASP 593 | - | PHE 594 | - | Yes | 0 | 0 | 0 | 0 | 0 | 0.55 | 0 | 0 | 1 | 3.01 |
| 32 | P194 | COC(=O)CNC(=O)Nc1ccc(cc1)Oc1ccnc(c1)C#N | 326.31 | 0.43 | -8.728 | -7.086 | ALA 480, LYS 482, LEU 513, THR 528 | LYS 482, GLU 500, CYS 531, ASP 593 | - | PHE 594 | - | Yes | 0 | 0 | 0 | 0 | 0 | 0.55 | 0 | 0 | 1 | 2.53 |
| 33 | P198 | O=C(Nc1ccc(c(c1)C(F)(F)F)Br)Nc1ccc(cc1)Oc1cnccc1C | 466.25 | 0.59 | -11.594 | -8.314 | ILE 462, VAL 470, LYS 482, LEU 504, LEU 513, THR 528, ASP 593 | LYS 482, GLU 500, CYS 531, ASP 593 | - | PHE 594 | - | Yes | 0 | 1 | 0 | 1 | 0 | 0.55 | 0 | 0 | 2 | 2.87 |
| 34 | P203 | COC(=O)c1nccc(c1)Oc1ccc(cc1)NC(=O)NCC1CC1 | 341.36 | 0.45 | -9.3 | -7.116 | ALA 480, LYS 482, GLU 500, VAL 503, LEU 504, THR 528 | LYS 482, GLU 500, CYS 531, ASP 593 | - | PHE 594 | - | Yes | 0 | 0 | 0 | 0 | 0 | 0.55 | 0 | 0 | 1 | 2.67 |
| 35 | P208 | COC(=O)c1nccc(c1)Oc1ccc(cc1)NC(=O)NC(=O)C(F)F | 365.29 | 0.48 | -9.759 | -8.351 | ALA 480, LYS 482, THR 528 | LYS 482, GLU 500, CYS 531, ASP 593 | - | PHE 594 | - | Yes | 0 | 0 | 0 | 0 | 0 | 0.55 | 0 | 0 | 2 | 2.59 |
| 36 | P214 | CCOC(=O)c1nccc(c1)Oc1ccc(cc1)NC(=O)NCC(=O)Nc1ccc(cc1)CN(C)C | 491.54 | 0.47 | -9.867 | -8.071 | LYS 482, VAL 503, THR 528, PHE 582, PHE 594 | LYS 482, GLU 500, CYS 531, ASP 593 | - | PHE 594 | - | Yes | 0 | 2 | 1 | 0 | 0 | 0.55 | 0 | 0 | 2 | 3.61 |
| 37 | P219 | O=NN(CCCN(c1ccc(cc1CN)NC(=O)Nc1ccc(cc1)Oc1cncc(c1)Cl)C)C | 497.98 | 0.42 | -9.552 | -7.315 | LYS 482, GLU 500, LEU 504, THR 528, PHE 594 | LYS 482, GLU 500, LEU 513, CYS 531, ILE 591, ASP 593 | - | PHE 594 | - | Yes | 0 | 2 | 1 | 0 | 0 | 0.55 | 1 | 1 | 2 | 3.59 |
| 38 | P220 | O=CCCNC(=O)Nc1ccc(cc1)Oc1ccnc(c1)C(Nc1ccccc1)OC | 420.46 | 0.42 | -9.535 | -8.238 | ALA 480, LYS 482, GLU 500, LEU 504, LEU 513, THR 528, PHE 582, PHE 594 | LYS 482, GLU 500, CYS 531, ASP 593 | - | PHE 594 | - | Yes | 0 | 0 | 1 | 0 | 0 | 0.55 | 0 | 2 | 2 | 3.68 |
| 39 | P238 | CNCCCN(c1ccc(cc1C)NC(=O)Nc1ccc(cc1)Oc1cncc(c1)C(=O)OC)C=O | 491.54 | 0.44 | -9.988 | -7.539 | ALA 480, GLU 500, VAL 503, LEU 504, THR 528, PHE 594 | LYS 482, GLU 500, CYS 531, HIS 573, ASP 593 | - | TRP 530, PHE 594 | - | Yes | 0 | 2 | 1 | 0 | 0 | 0.55 | 0 | 1 | 2 | 3.61 |
| 40 | P245 | N#CC(c1ccc(cc1)NC(=O)Nc1ccc(cc1)Oc1cnccc1C)(C(=O)O)C | 416.43 | 0.43 | -9.869 | -7.804 | ILE 462, VAL 470, ALA 480, GLU 500, VAL 503, LEU 504, THR 528, LEU 566, ASP 593, PHE 594 | LYS 482, GLU 500, CYS 531, ASP 593 | HIS 573 | TRP 530, PHE 594 | - | Yes | 0 | 0 | 0 | 0 | 0 | 0.56 | 0 | 0 | 2 | 3.46 |
| 41 | P248 | N#Cc1nccc(c1)Oc1ccc(cc1)NC(=O)NC(=O)C1CC1 | 322.32 | 0.41 | -9.828 | -7.875 | ALA 480, GLU 500, VAL 503, LEU 504, LEU 513, THR 528, ASP 593, PHE 594 | LYS 482, GLU 500, CYS 531, ASP 593 | - | PHE 594 | - | Yes | 0 | 0 | 0 | 0 | 0 | 0.55 | 0 | 0 | 0 | 2.43 |
| 42 | P251 | O=C(Nc1ccc(cc1)F)Nc1ccc(cc1)Oc1cncc(c1)Cl | 357.77 | 0.49 | -10.269 | -7.611 | ALA 480, LYS 482, GLU 500, VAL 503, LEU 504, THR 528 | LYS 482, GLU 500, THR 528, CYS 531, ASP 593 | - | PHE 594 | - | Yes | 0 | 0 | 0 | 0 | 0 | 0.55 | 0 | 0 | 2 | 2.66 |
| 43 | P268 | CCN(c1nccc(c1)Oc1ccc(cc1)NC(=O)NC(=O)OC(Br)C)C | 437.29 | 0.4 | -9.351 | -7.292 | ALA 480, LYS 482, LEU 513, THR 528, PHE 582, ASP 593, PHE 594 | LYS 482, GLU 500, CYS 531, ASP 593 | - | PHE 594 | - | Yes | 0 | 0 | 0 | 0 | 0 | 0.55 | 0 | 1 | 3 | 3.74 |
| 44 | P269 | CNc1ccc(cc1Br)NC(=O)Nc1ccc(cc1)Oc1cncc(c1)Cl | 447.71 | 0.52 | -10.367 | -8.003 | LYS 482, GLU 500, LEU 504, LEU 513, THR 528, ASP 593 | LYS 482, GLU 500, CYS 531, ASP 593 | - | PHE 594 | - | Yes | 0 | 0 | 0 | 0 | 0 | 0.55 | 0 | 0 | 2 | 3.08 |
| 45 | P280 | COC(=O)c1cncc(c1)Oc1ccc(cc1)NC(=O)Nc1ccc(cc1)Cl | 397.81 | 0.5 | -10.308 | -7.736 | ALA 480, LYS 482, GLU 500, VAL 503, LEU 504, THR 528 | LYS 482, GLU 500, CYS 531, ASP 593 | - | PHE 594 | - | Yes | 0 | 0 | 0 | 0 | 0 | 0.55 | 0 | 0 | 2 | 2.87 |
| 46 | P287 | O=C(Nc1ccc(c(c1)c1ccnc(c1)C)N(C)C)Nc1ccc(cc1)Oc1cccnc1 | 439.51 | 0.45 | -11.102 | -7.894 | ALA 480, ASN 499, GLU 500, VAL 503, LEU 504, THR 528, ASP 593, PHE 594 | LYS 482, GLU 500, CYS 531, ASP 593 | - | TRP 530, PHE 594 | - | Yes | 0 | 1 | 0 | 0 | 0 | 0.55 | 1 | 0 | 3 | 3.39 |
| 47 | P289 | COC(=O)CNC(=O)Nc1ccc(cc1)Oc1ccnc(c1)CN | 330.34 | 0.43 | -8.522 | -7.088 | LYS 482, LEU 513, THR 528 | LYS 482, GLU 500, CYS 531, ASP 593 | HIS 573 | PHE 594 | - | Yes | 0 | 0 | 0 | 0 | 0 | 0.55 | 0 | 0 | 1 | 2.5 |
| 48 | P303 | N#Cc1nccc(c1)Oc1ccc(cc1)NC(=O)NC(OS(=O)(=O)c1ccccc1C)C | 452.48 | 0.41 | -10.41 | -8.288 | ALA 480, LYS 482, VAL 503, LEU 504, LEU 513, THR 528, LEU 566, ILE 571, ASP 593 | LYS 482, GLU 500, CYS 531, ASP 593 | - | PHE 594 | - | Yes | 0 | 0 | 0 | 1 | 0 | 0.55 | 0 | 2 | 2 | 3.93 |
| 49 | P312 | NCc1nccc(c1)Oc1ccc(cc1)NC(=O)NC(=O)Cc1ccccc1 | 376.41 | 0.44 | -10.193 | -7.808 | ALA 480, LYS 482, GLU 500, VAL 503, LEU 504, THR 528, TRP 530, LEU 566, PHE 594 | LYS 482, GLU 500, CYS 531, ASP 593 | - | HIS 573, PHE 594 | - | Yes | 0 | 0 | 0 | 0 | 0 | 0.55 | 0 | 0 | 2 | 2.66 |
| 50 | P316 | ClCc1ccc(cc1)NC(=O)Nc1ccc(cc1)Oc1cncc(c1)F | 371.79 | 0.46 | -10.175 | -7.511 | ALA 480, LYS 482, GLU 500, VAL 503, LEU 504, THR 528 | LYS 482, GLU 500, CYS 531, ASP 593 | - | PHE 594 | - | Yes | 0 | 0 | 0 | 0 | 0 | 0.55 | 0 | 1 | 1 | 2.69 |
| 51 | P342 | N#Cc1nccc(c1)Oc1ccc(cc1)NC(=O)NC(OS(=O)(=O)c1ccccc1C(=O)O)C | 482.47 | 0.42 | -10.557 | -7.919 | ALA 480, LYS 482, VAL 503, LEU 504, LEU 513, THR 528, LEU 566, ILE 571, PHE 594 | LYS 482, GLU 500, CYS 531, ASP 593 | HIS 573 | - | - | Yes | 1 | 1 | 1 | 1 | 1 | 0.11 | 0 | 2 | 2 | 4.01 |
| 52 | P351 | N#CNc1ccc(cc1)Oc1ccnc(c1)CNC(=O)c1ccccc1NC | 373.41 | 0.42 | -8.828 | -8.001 | ALA 480, LYS 482, LEU 513, THR 528 | LYS 482, GLU 500, CYS 531, SER 535, ASP 593 | - | PHE 594 | - | Yes | 0 | 0 | 0 | 0 | 0 | 0.55 | 0 | 2 | 3 | 2.75 |
| 53 | P361 | NCc1nccc(c1)Oc1ccc(cc1)NC(=O)NC(=O)c1ccccc1Cl | 396.83 | 0.51 | -10.083 | -7.976 | LYS 482, GLU 500, VAL 503, LEU 513, THR 528, ASP 593 | LYS 482, GLU 500, CYS 531, ASP 593 | - | PHE 594 | - | Yes | 0 | 0 | 0 | 0 | 0 | 0.55 | 0 | 0 | 2 | 2.63 |
| 54 | P363 | CNCCCN(c1ccc(cc1C(=O)N)NC(=O)Nc1ccc(cc1)Oc1cncc(c1)Cl)C | 482.96 | 0.48 | -10.077 | -7.547 | LYS 482, GLU 500, VAL 503, LEU 504, THR 528 | LYS 482, GLU 500, LEU 513, CYS 531, HIS 573, ILE 591, ASP 593 | - | PHE 594 | - | Yes | 0 | 2 | 1 | 0 | 0 | 0.55 | 1 | 0 | 2 | 3.49 |
| 55 | P375 | CNCCCN(c1ccc(cc1C#N)NC(=O)Nc1ccc(cc1)Oc1cncc(c1)Cl)C=O | 478.93 | 0.45 | -10.015 | -7.59 | LYS 482, GLU 500, LEU 504, LEU 513, THR 528, ASP 593 | LYS 482, GLU 500, CYS 531, ASP 593 | - | PHE 594 | - | Yes | 0 | 1 | 1 | 0 | 0 | 0.55 | 0 | 1 | 2 | 3.39 |
| 56 | P419 | N#CC1(CC1)c1ccc(cc1)NC(=O)Nc1ccc(cc1)Oc1cccnc1 | 370.4 | 0.4 | -10.024 | -7.877 | ALA 480, GLU 500, VAL 503, LEU 504, THR 528, LEU 566, PHE 582, ASP 593, PHE 594 | LYS 482, GLU 500, CYS 531, ASP 593 | - | TRP 530, PHE 594 | - | Yes | 0 | 0 | 0 | 0 | 0 | 0.55 | 0 | 0 | 1 | 2.72 |

Supplementary Table 1: Molecule descriptors, Tanimoto similarity value, docking scores, target-ligand interactions, and ADME properties of 214 PURE ligands

| S. No. | Ligand | Canonical SMILES | Molecular weight (Da) | Tanimoto Similarity | Docking score with BRAF (kcal/mol) | Docking score with ABCG2 (kcal/mol) | Hydrophobic interactions | Hydrogen bonds | Salt bridge | Pi stack interaction(s) | Pi-cation interaction | Hydrogen bond match | Lipinski violation(s) | Ghose violation(s) | Veber violation(s) | Egan violation(s) | Muegge violation(s) | Bioavailability score | PAINS alert(s) | Brenk alert(s) | Leadlikeness violation(s) | Synthetic accessibility score |
|---|---|---|---|---|---|---|---|---|---|---|---|---|---|---|---|---|---|---|---|---|---|---|
| 57 | P423 | N=CCCNC(=O)Nc1ccc(cc1)Oc1ccnc(c1)C(=O)N1CCCC1 | 381.43 | 0.4 | -9.73 | -8.213 | ALA 480, LYS 482, THR 528, ASP 593 | LYS 482, GLU 500, CYS 531, ASP 593 | - | PHE 594 | - | Yes | 0 | 0 | 0 | 0 | 0 | 0.55 | 0 | 1 | 2 | 2.85 |
| 58 | P432 | CN(CCCN(c1ccc(cc1C(F)(F)F)NC(=O)Nc1ccc(cc1)Oc1cccnc1)C)C | 487.52 | 0.52 | -9.989 | -7.645 | GLU 500, LEU 504, THR 528, TRP 530, ASP 593 | LYS 482, GLU 500, CYS 531, ASP 593 | - | PHE 594 | - | Yes | 0 | 3 | 1 | 1 | 0 | 0.55 | 1 | 0 | 3 | 3.58 |
| 59 | P442 | N#Cc1nccc(c1)Oc1ccc(cc1)NC(=O)NC(=O)Cc1ccccc1O | 388.38 | 0.45 | -11.024 | -7.879 | ALA 480, LYS 482, VAL 503, LEU 504, LEU 513, THR 528, LEU 566, ILE 571 | LYS 482, GLU 500, CYS 531, HIS 573, ASP 593 | - | PHE 594 | - | Yes | 0 | 0 | 0 | 0 | 0 | 0.55 | 0 | 0 | 2 | 2.78 |
| 60 | P449 | CCOC(=O)CCCc1ccc(cc1)NC(=O)Nc1ccc(cc1)Oc1cncc(c1)F | 437.46 | 0.44 | -10.103 | -7.852 | ALA 480, LYS 482, GLU 500, VAL 503, LEU 504, THR 528, TRP 530, LEU 566, ILE 571 | LYS 482, GLU 500, CYS 531, ASP 593 | HIS 573 | PHE 594 | - | Yes | 0 | 0 | 1 | 0 | 0 | 0.55 | 0 | 0 | 3 | 3.17 |
| 61 | P455 | NC(=O)Nc1ccc(cc1)Oc1ccnc(c1)C(=O)N(C)C | 300.31 | 0.46 | -8.921 | -7.804 | ALA 480, LYS 482, LEU 513, THR 528 | LYS 482, GLU 500, CYS 531, ASP 593 | - | PHE 594 | - | Yes | 0 | 0 | 0 | 0 | 0 | 0.55 | 0 | 0 | 0 | 2.25 |
| 62 | P465 | N#Cc1nccc(c1)Oc1ccc(cc1)NC(=O)NC(=O)Cc1ccc(c(c1)C)N | 401.42 | 0.47 | -10.779 | -8.203 | ALA 480, LYS 482, GLU 500, VAL 503, LEU 504, THR 507, THR 528, TRP 530, LEU 566, ASP 593 | LYS 482, GLU 500, CYS 531, ASP 593 | - | PHE 594 | - | Yes | 0 | 0 | 0 | 0 | 0 | 0.55 | 0 | 1 | 2 | 2.9 |
| 63 | P473 | O=C(Nc1ccc(cc1)C1(CC1)C(=O)O)Nc1ccc(cc1)Oc1cncc(c1)Cl | 423.85 | 0.46 | -10.428 | -8.123 | ALA 480, GLU 500, VAL 503, LEU 504, THR 507, THR 528, TRP 530, LEU 566, ASP 593, PHE 594 | LYS 482, GLU 500, CYS 531, ASP 593 | HIS 573 | PHE 594 | - | Yes | 0 | 0 | 0 | 0 | 0 | 0.56 | 0 | 0 | 2 | 2.91 |
| 64 | P482 | NCCCNc1cc(ccc1OC)CC(=O)NC(=O)Nc1ccc(cc1)Oc1ccnc(c1)C#N | 474.51 | 0.43 | -8.698 | -7.619 | LYS 482, GLU 500, VAL 503, LEU 513, THR 528, ASP 593 | LYS 482, ALA 496, GLU 500, CYS 531, ASP 593 | - | PHE 594 | - | Yes | 0 | 1 | 2 | 1 | 1 | 0.55 | 0 | 0 | 2 | 3.47 |
| 65 | P483 | NCc1nccc(c1)Oc1ccc(cc1)NC(=O)NC(=O)OC(=O)C | 344.32 | 0.42 | -8.434 | -7.508 | VAL 470, LYS 482, VAL 503, LEU 504, TRP 530, PHE 594 | LYS 482, GLU 500, CYS 531, ASP 593 | - | PHE 594 | - | Yes | 0 | 0 | 0 | 1 | 0 | 0.55 | 0 | 2 | 1 | 2.63 |
| 66 | P489 | O=NN(c1ccccc1C(=O)NC(=O)Nc1ccc(cc1)Oc1ccnc(c1)C(=O)NC(=O)C)C | 476.44 | 0.47 | -10.945 | -8.43 | ALA 480, LYS 482, GLU 500, VAL 503, LEU 513, THR 528, ASP 593 | LYS 482, GLU 500, CYS 531, ASP 593 | - | PHE 594 | - | Yes | 1 | 0 | 2 | 1 | 1 | 0.55 | 0 | 1 | 2 | 3.2 |
| 67 | P498 | CCN(c1nccc(c1)Oc1ccc(cc1)NC(=O)NC(=O)OC(=O)C)C | 372.38 | 0.4 | -9.452 | -7.418 | ALA 480, LYS 482, THR 528, TRP 530, ASP 593 | LYS 482, GLU 500, CYS 531, ASP 593 | - | PHE 594 | - | Yes | 0 | 0 | 0 | 0 | 0 | 0.55 | 0 | 2 | 2 | 3.08 |
| 68 | P499 | O=C(Nc1ccc(c(c1)Oc1cncc(c1Cl)F)N(C)C)Nc1ccc(cc1)Oc1ccncc1 | 493.92 | 0.44 | -11.11 | -8.372 | ALA 480, LYS 482, GLU 500, LEU 504, THR 528 | LYS 482, GLU 500, CYS 531, ASP 593 | - | PHE 594 | - | Yes | 0 | 3 | 0 | 1 | 0 | 0.55 | 1 | 0 | 3 | 3.62 |
| 69 | P515 | CNCCCN(c1ccc(cc1CN)NC(=O)Nc1ccc(cc1)Oc1cncc(c1)Cl)C | 468.98 | 0.45 | -9.289 | -7.145 | VAL 470, GLU 500, VAL 503, LEU 504, LEU 513, THR 528, ILE 571 | LYS 482, GLU 500, CYS 531, ILE 572, ILE 591, ASP 593 | - | PHE 594 | - | Yes | 0 | 1 | 1 | 0 | 0 | 0.55 | 1 | 0 | 2 | 3.5 |
| 70 | P534 | NCCc1ccc(cc1)NC(=O)Nc1ccc(cc1)Oc1cncc(c1)C(=O)C=NO | 419.43 | 0.42 | -9.765 | -8.034 | LYS 482, GLU 500, VAL 503, LEU 504, THR 528 | LYS 482, GLU 500, CYS 531, ASP 593 | - | PHE 594 | - | Yes | 0 | 0 | 0 | 1 | 0 | 0.55 | 0 | 3 | 2 | 3.2 |
| 71 | P537 | COC(=O)c1cncc(c1)Oc1ccc(cc1)NC(=O)Nc1ccc(c(c1)OC(F)(F)F)NC | 476.41 | 0.55 | -11 | -8.24 | LYS 482, GLU 500, VAL 503, LEU 504, LEU 513, THR 528, ASP 593 | LYS 482, GLU 500, CYS 531, ASP 593 | - | PHE 594 | - | Yes | 0 | 1 | 1 | 1 | 0 | 0.55 | 0 | 0 | 3 | 3.41 |
| 72 | P538 | N#CC1(CC1)c1ccc(cc1)NC(=O)Nc1ccc(cc1)Oc1cncc(c1)F | 388.39 | 0.42 | -10.525 | -7.882 | LYS 482, GLU 500, VAL 503, LEU 504, THR 507, ILE 512, THR 528, LEU 566, ASP 593, PHE 594 | LYS 482, GLU 500, CYS 531, ASP 593 | - | TRP 530, PHE 594 | - | Yes | 0 | 0 | 0 | 0 | 0 | 0.55 | 0 | 0 | 1 | 2.83 |
| 73 | P540 | N#Cc1nccc(c1)Oc1ccc(cc1)NC(=O)NCC(=O)Nc1ccc(cc1Br)F | 484.28 | 0.49 | -10.628 | -8.331 | LYS 482, VAL 503, LEU 513, THR 528, ILE 571 | LYS 482, GLU 500, CYS 531, ASP 593 | - | PHE 594 | - | Yes | 0 | 1 | 0 | 0 | 0 | 0.55 | 0 | 0 | 2 | 2.93 |
| 74 | P548 | CC(OC(=O)NC(=O)Nc1ccc(cc1)Oc1ccnc(c1)C(=O)N(C)C)Cl | 406.82 | 0.44 | -9.43 | -7.598 | ALA 480, LYS 482, VAL 503, LEU 504, LEU 513, THR 528 | LYS 482, GLU 500, CYS 531, ASP 593 | - | PHE 594 | - | Yes | 0 | 0 | 0 | 0 | 0 | 0.55 | 0 | 1 | 2 | 3.38 |
| 75 | P554 | C=COC(=O)Nc1ccc(c(c1)F)Oc1ccnc(c1)C(=O)NC | 331.3 | 0.44 | -9.15 | -7.462 | LYS 482, LEU 504, LEU 513, THR 528 | LYS 482, GLU 500, CYS 531, ASP 593 | - | PHE 594 | - | Yes | 0 | 0 | 0 | 0 | 0 | 0.55 | 0 | 1 | 1 | 2.64 |
| 76 | P559 | CNc1cccc(c1)NC(=O)Nc1ccc(cc1)NC(=O)c1cncc(c1)C(Cl)(Cl)Cl | 478.76 | 0.42 | -11.085 | -8.314 | LYS 482, GLU 500, VAL 503, LEU 504, LEU 513, THR 528, ASP 593, PHE 594 | LYS 482, GLU 500, CYS 531, ASP 593 | - | TRP 530, PHE 594 | - | Yes | 0 | 0 | 0 | 0 | 0 | 0.55 | 0 | 1 | 3 | 2.86 |
| 77 | P569 | O=C(Nc1ccc(c(c1)Cl)N(C)C)Nc1ccc(cc1)Oc1ccncc1 | 382.84 | 0.52 | -10.144 | -7.325 | ALA 480, LYS 482, GLU 500, LEU 504, THR 528, ASP 593, PHE 594 | LYS 482, GLU 500, CYS 531, ASP 593 | - | PHE 594 | - | Yes | 0 | 0 | 0 | 0 | 0 | 0.55 | 1 | 0 | 2 | 2.64 |
| 78 | P607 | O=C(NC(=O)c1ccccc1F)Nc1ccc(cc1)Oc1ccnc(c1)C(=O)C | 393.37 | 0.53 | -11.235 | -8.428 | LYS 482, GLU 500, VAL 503, LEU 504, LEU 513, THR 528, TRP 530, PHE 582, ASP 593 | LYS 482, GLU 500, CYS 531, ASP 593 | - | PHE 594 | - | Yes | 0 | 0 | 0 | 0 | 0 | 0.55 | 0 | 0 | 2 | 2.7 |
| 79 | P622 | CCOc1cc(ccc1C)NC(=O)Nc1ccc(cc1)Oc1cccnc1 | 363.41 | 0.46 | -9.792 | -7.526 | GLU 500, VAL 503, LEU 504, THR 528, ASP 593, PHE 594 | LYS 482, GLU 500, CYS 531, ASP 593 | - | PHE 594 | - | Yes | 0 | 0 | 0 | 0 | 0 | 0.55 | 0 | 0 | 3 | 2.96 |
| 80 | P625 | COC(=O)c1nccc(c1)Oc1ccc(cc1)NC(=O)NC(=O)C1CC1 | 355.34 | 0.45 | -9.803 | -7.931 | ALA 480, LYS 482, GLU 500, VAL 503, LEU 504, THR 528, ASP 593 | LYS 482, GLU 500, CYS 531, ASP 593 | - | PHE 594 | - | Yes | 0 | 0 | 0 | 0 | 0 | 0.55 | 0 | 0 | 2 | 2.64 |
| 81 | P639 | O=CNNc1ccc(cc1)Oc1ccnc(c1)CNC(=O)c1ccccc1C(F)F | 412.39 | 0.41 | -9.545 | -8.41 | ILE 462, ALA 480, LYS 482, THR 528 | LYS 482, GLU 500, CYS 531, ASP 593 | - | TRP 530, PHE 594 | - | Yes | 0 | 0 | 0 | 0 | 0 | 0.55 | 0 | 1 | 2 | 2.84 |
| 82 | P649 | NCc1nccc(c1)Oc1ccc(cc1)NC(=O)NC(=O)c1ccccc1NCCC(=O)O | 449.46 | 0.43 | -10.369 | -7.934 | LYS 482, GLU 500, VAL 503, LEU 513, THR 528, LEU 566, ASP 593 | LYS 482, GLU 500, LEU 513, CYS 531, ASP 593 | - | PHE 594 | - | Yes | 0 | 0 | 2 | 1 | 1 | 0.55 | 0 | 0 | 2 | 3.05 |
| 83 | P666 | O=C(Nc1ccc(cc1)[N+](=O)[O-])NCc1ccc(cc1)Oc1ccnc(c1)C(N)C | 407.42 | 0.43 | -9.964 | -8.285 | VAL 470, GLU 500, VAL 503, LEU 504, PHE 582, PHE 594 | LYS 482, GLU 500, CYS 531, ASP 593 | - | - | - | Yes | 0 | 0 | 0 | 1 | 0 | 0.55 | 0 | 2 | 2 | 3.38 |
| 84 | P677 | O=C(Nc1ccc(cc1)Oc1ccnc(c1)CN1CCCC1)NC(=O)C | 354.4 | 0.4 | -9.291 | -8.007 | ALA 480, LYS 482, GLU 500, THR 528, TRP 530, PHE 582, ASP 593, PHE 594 | LYS 482, GLU 500, CYS 531, ASP 593 | - | PHE 594 | - | Yes | 0 | 0 | 0 | 0 | 0 | 0.55 | 0 | 0 | 2 | 2.6 |

#### Supplementary Table 1: Molecule descriptors, Tanimoto similarity value, docking scores, target-ligand interactions, and ADME properties of 214 PURE ligands

| S. No. | Ligand | Canonical SMILES | Molecular weight (Da) | Tanimoto Similarity | Docking score with BRAF (kcal/mol) | Docking score with ABCG2 (kcal/mol) | Hydrophobic interactions | Hydrogen bonds | Salt bridge | Pi stack interaction(s) | Pi-cation interaction | Hydrogen bond match | Lipinski violation(s) | Ghose violation(s) | Veber violation(s) | Egan violation(s) | Muegge violation(s) | Bioavailability score | PAINS alert(s) | Brenk alert(s) | Leadlikeness violation(s) | Synthetic accessibility score |
|---|---|---|---|---|---|---|---|---|---|---|---|---|---|---|---|---|---|---|---|---|---|---|
| 85 | P682 | CCOC(=O)Nc1ccc(c(c1)F)Oc1ccnc(c1)C(=O)NC | 333.31 | 0.44 | -9.212 | -7.561 | ALA 480, LYS 482, GLU 500, LEU 504, LEU 513, THR 528 | LYS 482, GLU 500, CYS 531, ASP 593 | - | PHE 594 | - | Yes | 0 | 0 | 0 | 0 | 0 | 0.55 | 0 | 0 | 1 | 2.65 |
| 86 | P686 | CCC(=O)c1nccc(c1)Oc1ccc(cc1)NC(=O)NCC(=O)Nc1ccc(cc1)C(N(C)C)C | 489.57 | 0.47 | -10.394 | -8.39 | ALA 480, LYS 482, GLU 500, VAL 503, LEU 513, THR 528, TRP 530, PHE 582 | LYS 482, GLU 500, CYS 531, ASP 593 | - | PHE 594 | - | Yes | 0 | 2 | 1 | 0 | 0 | 0.55 | 0 | 0 | 2 | 4 |
| 87 | P687 | CNC(=O)Nc1ccc(cc1)Oc1ccnc(c1)C(=O)C | 285.3 | 0.54 | -9.055 | -7.584 | ALA 480, LYS 482, THR 528, PHE 582 | LYS 482, GLU 500, CYS 531, ASP 593 | - | PHE 594 | - | Yes | 0 | 0 | 0 | 0 | 0 | 0.55 | 0 | 0 | 0 | 2.25 |
| 88 | P703 | CON(C(=O)c1cncc(c1)Oc1ccc(cc1)NC(=O)Nc1ccc(cc1)N(N=O)C)C | 450.45 | 0.42 | -10.228 | -8.226 | LYS 482, GLU 500, LEU 504, THR 528, ASP 593 | LYS 482, GLU 500, CYS 531, HIS 573, ASP 593 | - | TRP 530, PHE 594 | - | Yes | 1 | 0 | 1 | 0 | 0 | 0.55 | 0 | 2 | 2 | 3.45 |
| 89 | P708 | O=C(OC(C)(C)C)NC(=O)Nc1ccc(cc1)Oc1ccnc(c1)C(=O)N(C)C | 400.43 | 0.44 | -10.276 | -7.766 | ALA 480, LEU 504, LEU 513, THR 528, ASP 593 | LYS 482, GLU 500, CYS 531, ASP 593 | - | PHE 594 | - | Yes | 0 | 0 | 0 | 0 | 0 | 0.55 | 0 | 0 | 2 | 3.03 |
| 90 | P709 | COC(=O)c1nccc(c1)Oc1ccc(cc1)NC(=O)NC(=O)OCCCC(O)(C)C | 431.44 | 0.41 | -8.656 | -7.048 | ALA 480, VAL 503, LEU 504, THR 528, TRP 530, LEU 566, PHE 594 | LYS 482, GLU 500, CYS 531, ASP 593 | - | PHE 594 | - | Yes | 0 | 0 | 1 | 1 | 0 | 0.55 | 0 | 1 | 2 | 3.45 |
| 91 | P720 | CNc1ccccc1C(=O)NC(=O)Nc1ccc(cc1)Oc1ccnc(c1)C(=O)C | 404.42 | 0.53 | -10.917 | -7.9 | ALA 480, LYS 482, GLU 500, VAL 503, LEU 504, THR 528, TRP 530, PHE 582, ASP 593 | LYS 482, GLU 500, CYS 531, HIS 573, ASP 593 | - | PHE 594 | - | Yes | 0 | 0 | 0 | 0 | 0 | 0.55 | 0 | 0 | 2 | 2.85 |
| 92 | P733 | N#Cc1nccc(c1)Oc1ccc(cc1)NC(=O)NC(=O)OC(C)(C)C | 354.36 | 0.42 | -9.461 | -7.172 | LYS 482, LEU 504, LEU 513, THR 528, ASP 593 | LYS 482, GLU 500, CYS 531, ASP 593 | - | PHE 594 | - | Yes | 0 | 0 | 0 | 0 | 0 | 0.55 | 0 | 0 | 2 | 2.87 |
| 93 | P734 | O=C(NC(=O)c1ccccc1Cl)Nc1ccc(cc1)Oc1ccnc(c1)C(=O)C | 409.82 | 0.55 | -10.984 | -8.341 | ALA 480, GLU 500, VAL 503, LEU 513, THR 528, TRP 530, PHE 582 | LYS 482, GLU 500, CYS 531, ASP 593 | - | PHE 594 | - | Yes | 0 | 0 | 0 | 0 | 0 | 0.55 | 0 | 0 | 3 | 2.69 |
| 94 | P735 | O=C(Nc1ccc(c(c1)C(c1ccc(cn1)O)O)N(C)C)Nc1ccc(cc1)Oc1ccncc1 | 471.51 | 0.42 | -10.657 | -8.414 | ALA 480, LYS 482, VAL 503, LEU 504, LEU 513, THR 528, ASP 593 | LYS 482, GLU 500, CYS 531, HIS 573, ASP 593 | - | PHE 594 | - | Yes | 0 | 1 | 0 | 0 | 0 | 0.55 | 1 | 0 | 2 | 3.9 |
| 95 | P737 | NCC(c1ccc(cc1)NC(=O)Nc1ccc(cc1)Oc1cncc(c1)C(=O)OC)C(=O)O | 450.44 | 0.42 | -9.451 | -7.755 | ALA 480, GLU 500, VAL 503, LEU 504, THR 528, PHE 594 | LYS 482, GLU 500, CYS 531, HIS 573, ASP 593 | - | TRP 530, PHE 594 | - | Yes | 0 | 0 | 2 | 1 | 1 | 0.55 | 0 | 0 | 2 | 3.68 |
| 96 | P740 | COC(=O)c1nccc(c1)Oc1ccc(cc1)NC(=O)NC(=O)OC(Br)C | 438.23 | 0.44 | -9.081 | -7.185 | ALA 480, GLU 500, VAL 503, LEU 504, THR 528 | LYS 482, GLU 500, CYS 531, ASP 593 | - | PHE 594 | - | Yes | 0 | 0 | 0 | 0 | 0 | 0.55 | 0 | 2 | 3 | 3.49 |
| 97 | P741 | O=C(Nc1ccc(cc1)Oc1ccnc(c1)C(=O)N(C)C)NCc1ccccc1 | 390.44 | 0.48 | -10.609 | -8.035 | ALA 480, LYS 482, THR 528, ASP 593 | LYS 482, GLU 500, CYS 531, ASP 593 | - | PHE 594 | - | Yes | 0 | 0 | 0 | 0 | 0 | 0.55 | 0 | 0 | 2 | 2.81 |
| 98 | P756 | CC(=O)CNC(=O)Nc1ccc(cc1)Oc1ccnc(c1)N(C)C | 328.37 | 0.43 | -9.027 | -7.207 | ALA 480, LYS 482, GLU 500, VAL 503, LEU 504, THR 528 | LYS 482, GLU 500, CYS 531, ASP 593 | - | PHE 594 | - | Yes | 0 | 0 | 0 | 0 | 0 | 0.55 | 0 | 0 | 1 | 2.68 |
| 99 | P771 | NCc1cc(ccc1N(CCCN(C)C)C)NC(=O)Nc1ccc(cc1)Oc1ccncc1 | 448.56 | 0.42 | -9.347 | -7.318 | ALA 480, LYS 482, GLU 500, VAL 503, LEU 504, LEU 513, THR 528 | LYS 482, GLU 500, CYS 531, GLY 592, ASP 593 | - | PHE 594 | - | Yes | 0 | 1 | 1 | 0 | 0 | 0.55 | 1 | 0 | 2 | 3.3 |
| 100 | P776 | CCC(c1nccc(c1)Oc1ccc(cc1)NC(=O)NCC(=O)Nc1ccc(cc1)CN(C)C)(O)C | 491.58 | 0.45 | -10.497 | -7.85 | ALA 480, LYS 482, VAL 503, LEU 513, THR 528, TRP 530, ILE 571, PHE 582, PHE 594 | LYS 482, GLU 500, CYS 531, ILE 572, ASP 593 | - | PHE 594 | - | Yes | 0 | 2 | 1 | 0 | 0 | 0.55 | 0 | 0 | 2 | 4.1 |
| 101 | P784 | CNc1cccc(c1)NC(=O)Nc1ccc(cc1)NC(=O)c1cncc(c1)C(F)(F)F | 429.4 | 0.49 | -10.156 | -8.249 | GLU 500, VAL 503, LEU 504, LEU 513, THR 528, ASP 593, PHE 594 | LYS 482, GLU 500, CYS 531, ASP 593 | - | TRP 530, PHE 594 | - | Yes | 0 | 0 | 0 | 0 | 0 | 0.55 | 0 | 0 | 2 | 2.92 |
| 102 | P788 | COC(=O)c1cncc(c1)Oc1ccc(cc1)NC(=O)Nc1ccc(cc1)C(=O)C | 405.4 | 0.45 | -10.268 | -7.809 | LYS 482, GLU 500, VAL 503, LEU 504, THR 528, LEU 566, ILE 571 | LYS 482, GLU 500, CYS 531, ASP 593 | - | PHE 594 | - | Yes | 0 | 0 | 0 | 0 | 0 | 0.55 | 0 | 0 | 2 | 2.98 |
| 103 | P799 | COc1ccc(cc1)NC(=O)Nc1ccc(cc1)Oc1cncc(c1)Cl | 369.8 | 0.48 | -10.266 | -7.331 | LYS 482, GLU 500, VAL 503, LEU 504, LEU 513, THR 528 | LYS 482, GLU 500, CYS 531, ASP 593 | - | PHE 594 | - | Yes | 0 | 0 | 0 | 0 | 0 | 0.55 | 0 | 0 | 2 | 2.83 |
| 104 | P800 | CCOC(=O)NC(=O)Nc1ccc(cc1)Oc1ccnc(c1)C(=O)C | 343.33 | 0.46 | -9.529 | -7.886 | ALA 480, LYS 482, VAL 503, THR 528 | LYS 482, GLU 500, CYS 531, ASP 593 | - | PHE 594 | - | Yes | 0 | 0 | 0 | 0 | 0 | 0.55 | 0 | 0 | 1 | 2.67 |
| 105 | P802 | C=COC(=O)N(CCCN(c1ccc(cc1C)NC(=O)Nc1ccc(cc1)Oc1cccnc1)C)C | 489.57 | 0.41 | -9.807 | -7.702 | LYS 482, GLU 500, VAL 503, LEU 504, THR 528, ILE 571 | LYS 482, GLU 500, CYS 531, ASP 593 | - | PHE 594 | - | Yes | 0 | 2 | 1 | 0 | 0 | 0.55 | 1 | 1 | 3 | 3.77 |
| 106 | P806 | C=COC(=O)NC(=O)Nc1ccc(cc1)Oc1ccnc(c1)C(=O)Cc1ccc(cc1)N(C)C | 460.48 | 0.45 | -10.617 | -8.145 | ILE 462, LYS 482, LEU 513, THR 528, TRP 530, PHE 594 | LYS 482, GLU 500, CYS 531, ASP 593 | - | PHE 582, PHE 594 | - | Yes | 0 | 0 | 1 | 0 | 0 | 0.55 | 1 | 1 | 3 | 3.53 |
| 107 | P813 | N#Cc1nccc(c1)Oc1ccc(cc1)NC(=O)NCC(=O)Nc1ccc(cc1)S(=O)(=O)Cl | 485.9 | 0.47 | -10.344 | -8.043 | ALA 480, LYS 482, VAL 503, LEU 513, THR 528, ILE 571 | LYS 482, GLU 500, CYS 531, ASP 593 | - | PHE 594 | - | Yes | 0 | 1 | 1 | 1 | 1 | 0.55 | 0 | 0 | 2 | 3.04 |
| 108 | P838 | CNC(=O)c1nccc(c1)Oc1ccc(cc1)NC(=O)NC(=O)OCCCCC(O)(C)C | 444.48 | 0.48 | -9.317 | -7.776 | LYS 482, VAL 503, LEU 504, THR 507, ILE 512, TRP 530, LEU 566, ILE 591, PHE 594 | LYS 482, GLU 500, CYS 531, ASP 593 | - | PHE 594 | - | Yes | 0 | 0 | 1 | 1 | 0 | 0.55 | 0 | 0 | 2 | 3.42 |
| 109 | P843 | CCOc1cc(ccc1OCC)NC(=O)Nc1ccc(cc1)Oc1cnccc1C | 407.46 | 0.42 | -10.04 | -7.587 | ILE 462, VAL 470, LYS 482, GLU 500, VAL 503, LEU 504, THR 528, LEU 566, ILE 571, ASP 593, PHE 594 | LYS 482, GLU 500, CYS 531, ASP 593 | - | TRP 530, PHE 594 | - | Yes | 0 | 0 | 0 | 0 | 0 | 0.55 | 0 | 0 | 3 | 3.34 |
| 110 | P850 | CNC(=O)c1nccc(c1)Oc1ccc(cc1)CNC(=O)Nc1ccc(cc1)[N+](=O)[O-] | 421.41 | 0.57 | -10.429 | -8.201 | VAL 470, LYS 482, GLU 500, LEU 504, LEU 513, PHE 594 | LYS 482, GLU 500, CYS 531, ASP 593 | - | - | - | Yes | 0 | 0 | 0 | 1 | 0 | 0.55 | 0 | 2 | 2 | 2.91 |

Supplementary Table 1: Molecule descriptors, Tanimoto similarity value, docking scores, target-ligand interactions, and ADME properties of 214 PURE ligands

| S. No. | Ligand | Canonical SMILES | Molecular weight (Da) | Tanimoto Similarity | Docking score with BRAF (kcal/mol) | Docking score with ABCG2 (kcal/mol) | Hydrophobic interactions | Hydrogen bonds | Salt bridge | Pi stack interaction(s) | Pi-cation interaction | Hydrogen bond match | Lipinski violation(s) | Ghose violation(s) | Veber violation(s) | Egan violation(s) | Muegge violation(s) | Bioavailability score | PAINS alert(s) | Brenk alert(s) | Leadlikeness violation(s) | Synthetic accessibility score |
|---|---|---|---|---|---|---|---|---|---|---|---|---|---|---|---|---|---|---|---|---|---|---|
| 111 | P851 | CCOc1ccc(cc1C(F)(F)F)NC(=O)Nc1ccc(cc1)Oc1cnccc1C(=O)O | 461.39 | 0.57 | -10.961 | -8.371 | LYS 482, GLU 500, VAL 503, LEU 504, LEU 513, THR 528, ILE 571 | LYS 482, GLU 500, CYS 531, ASP 593 | - | PHE 594 | - | Yes | 0 | 1 | 0 | 1 | 0 | 0.56 | 0 | 0 | 3 | 3.23 |
| 112 | P866 | NC(=O)Nc1ccc(cc1C(=N)Nc1ccc(cc1)Oc1ccncc1)NC(=O)C(SC(=O)C)C | 492.55 | 0.41 | -9.804 | -8.183 | VAL 470, ALA 480, LYS 482, GLU 500, LEU 504, LEU 513, THR 528, ASP 593, PHE 594 | LYS 482, GLU 500, CYS 531, ASP 593 | - | PHE 594 | LYS 482 | Yes | 0 | 2 | 2 | 1 | 1 | 0.55 | 0 | 3 | 2 | 3.9 |
| 113 | P899 | COc1cc(ccc1OC)CC(=O)NC(=O)Nc1ccc(cc1)Oc1ccnc(c1)C(=O)C | 449.46 | 0.48 | -9.933 | -8.154 | ILE 462, VAL 470, GLU 500, PHE 582, ASP 593, PHE 594 | LYS 482, GLU 500, CYS 531, ASP 593 | - | TRP 530, PHE 594 | - | Yes | 0 | 0 | 1 | 0 | 0 | 0.55 | 0 | 0 | 2 | 3.12 |
| 114 | P911 | CC(=O)OC(=O)NC(=O)Nc1ccc(cc1)Oc1ccnc(c1)C(=O)C | 357.32 | 0.45 | -8.874 | -7.462 | ALA 480, LYS 482, VAL 503, LEU 504, THR 528, TRP 530, PHE 582, PHE 594 | LYS 482, GLU 500, CYS 531, ASP 593 | HIS 573 | PHE 594 | - | Yes | 0 | 0 | 0 | 0 | 0 | 0.55 | 0 | 2 | 2 | 2.68 |
| 115 | P912 | O=C(NC(=O)c1ccccc1)Nc1ccc(cc1)Oc1ccnc(c1)C(=O)C | 375.38 | 0.51 | -10.964 | -8.271 | ILE 462, ALA 480, GLU 500, VAL 503, LEU 513, THR 528, ASP 593 | LYS 482, GLU 500, CYS 531, ASP 593 | - | PHE 594 | - | Yes | 0 | 0 | 0 | 0 | 0 | 0.55 | 0 | 0 | 2 | 2.57 |
| 116 | P931 | O=C(Nc1ccc(c(c1)C(Cl)(Cl)Cl)N)Nc1ccc(cc1)Oc1cccnc1 | 437.71 | 0.5 | -10.854 | -8.177 | LYS 482, GLU 500, LEU 504, LEU 513, THR 528, ASP 593 | LYS 482, GLU 500, CYS 531, ASP 593 | - | PHE 594 | - | Yes | 0 | 0 | 0 | 0 | 0 | 0.55 | 0 | 2 | 2 | 2.78 |
| 117 | P944 | CCOC(=O)NC(=O)Nc1ccc(c(c1)Oc1ccnc(c1)C(=O)NC)O | 374.35 | 0.44 | -9.319 | -7.989 | ALA 480, LYS 482, GLU 500, VAL 503, LEU 504, LEU 513, ILE 526, THR 528, PHE 594 | LYS 482, GLU 500, CYS 531, ASP 593 | - | - | - | Yes | 0 | 0 | 0 | 1 | 0 | 0.55 | 0 | 1 | 2 | 2.92 |
| 118 | P957 | O=C(Nc1ccc(cc1)Oc1ccnc(c1)C(=O)C)NCC1CC1 | 325.36 | 0.47 | -9.67 | -7.365 | ILE 462, ALA 480, LYS 482, GLU 500, VAL 503, LEU 504, LEU 513, THR 528 | LYS 482, GLU 500, CYS 531, ASP 593 | - | PHE 594 | - | Yes | 0 | 0 | 0 | 0 | 0 | 0.55 | 0 | 0 | 1 | 2.49 |
| 119 | P971 | CCC(c1ccc(cc1)NC(=O)Nc1ccc(cc1)Oc1cnccc1C)(C(=O)O)C | 419.47 | 0.43 | -10.226 | -7.896 | ILE 462, ALA 480, GLU 500, VAL 503, LEU 504, THR 507, THR 528, LEU 566, PHE 582, ASP 593, PHE 594 | LYS 482, GLU 500, CYS 531, ASP 593 | HIS 573 | TRP 530, PHE 594 | - | Yes | 0 | 0 | 0 | 0 | 0 | 0.56 | 0 | 0 | 3 | 3.55 |
| 120 | P982 | COC(=O)c1cncc(c1)Oc1ccc(cc1)NC(=O)Nc1ccc(cc1)N(CCCN(C)C)C | 477.56 | 0.4 | -9.739 | -7.315 | LYS 482, GLU 500, VAL 503, LEU 504, LEU 513, THR 528 | LYS 482, GLU 500, CYS 531, ASP 593 | - | PHE 594 | - | Yes | 0 | 1 | 1 | 0 | 0 | 0.55 | 1 | 0 | 2 | 3.62 |
| 121 | P985 | N#Cc1nccc(c1)Oc1ccc(cc1)NC(=O)NC(=O)OS(=O)(=O)c1cccc(c1C)O | 468.44 | 0.44 | -11.28 | -8.046 | ALA 480, LYS 482, VAL 503, LEU 513, THR 528, TRP 530, ILE 571 | LYS 482, GLU 500, CYS 531, ILE 572, ASP 593 | HIS 573 | PHE 594 | - | Yes | 1 | 0 | 1 | 1 | 1 | 0.55 | 0 | 1 | 2 | 3.41 |
| 122 | P994 | O=NN(c1ccc(cc1)NC(=O)Nc1ccc(cc1)Oc1cncc(c1)C(=O)N)C | 406.39 | 0.45 | -10.33 | -7.837 | LYS 482, GLU 500, VAL 503, LEU 504, THR 528 | LYS 482, GLU 500, CYS 531, ASP 593 | - | PHE 594 | - | Yes | 0 | 0 | 0 | 1 | 0 | 0.55 | 0 | 1 | 2 | 3.06 |
| 123 | P995 | O=C(Nc1ccc(cc1)Oc1ccnc(c1)C(=O)N(C)C)NC(=O)C | 342.35 | 0.46 | -9.82 | -7.813 | ALA 480, LYS 482, GLU 500, THR 528, ASP 593, PHE 594 | LYS 482, GLU 500, CYS 531, ASP 593 | - | PHE 594 | - | Yes | 0 | 0 | 0 | 0 | 0 | 0.55 | 0 | 0 | 1 | 2.5 |
| 124 | P998 | COC(=O)c1nccc(c1)Oc1ccc(cc1)NC(=O)NCC(=O)Nc1ccc(cc1)C(=O)N(C)C | 491.5 | 0.47 | -10.612 | -8.073 | ALA 480, LYS 482, VAL 503, LEU 513, THR 528 | LYS 482, GLU 500, CYS 531, ASP 593 | - | PHE 594 | - | Yes | 1 | 2 | 1 | 1 | 0 | 0.55 | 0 | 0 | 2 | 3.4 |
| 125 | P1002 | ClCc1nccc(c1)Oc1ccc(cc1)NC(=O)NC(=O)OC(=O)C | 363.75 | 0.43 | -8.997 | -7.541 | ALA 480, LYS 482, LEU 513, THR 528, ASP 593 | LYS 482, GLU 500, CYS 531, ASP 593 | - | PHE 594 | - | Yes | 0 | 0 | 0 | 0 | 0 | 0.55 | 0 | 3 | 2 | 2.72 |
| 126 | P1027 | C=COC(=O)N(CCCN(c1ccc(cc1C)NC(=O)Nc1ccc(cc1)Oc1ccncc1)C)C | 489.57 | 0.43 | -9.622 | -7.285 | ALA 480, LYS 482, GLU 500, VAL 503, LEU 504, LEU 513, THR 528, ILE 571 | LYS 482, GLU 500, CYS 531, ASP 593 | - | TRP 530, PHE 594 | - | Yes | 0 | 2 | 1 | 0 | 0 | 0.55 | 1 | 1 | 3 | 3.62 |
| 127 | P1038 | O=C(Nc1ccc(cc1)Oc1ccnc(c1)N(C)C)NC(=O)OC(Br)C | 423.26 | 0.41 | -8.647 | -6.984 | VAL 470, ALA 480, VAL 503, LEU 504, THR 528 | LYS 482, GLU 500, CYS 531, ASP 593 | - | TRP 530, PHE 594 | - | Yes | 0 | 0 | 0 | 0 | 0 | 0.55 | 0 | 1 | 3 | 3.65 |
| 128 | P1055 | COC(=O)c1nccc(c1)Oc1ccc(cc1)NC(=O)NC(=O)Cc1ccccc1F | 423.39 | 0.49 | -11.119 | -7.968 | ALA 480, LYS 482, VAL 503, THR 528, ILE 571, ASP 593 | LYS 482, GLU 500, CYS 531, ASP 593 | - | PHE 594 | - | Yes | 0 | 0 | 0 | 0 | 0 | 0.55 | 0 | 0 | 3 | 2.98 |
| 129 | P1070 | N#Cc1nccc(c1)Oc1ccc(cc1)NC(=O)NC(=O)C | 296.28 | 0.45 | -9.434 | -7.677 | ALA 480, LYS 482, THR 528, ASP 593, PHE 594 | LYS 482, GLU 500, CYS 531, ASP 593 | - | PHE 594 | - | Yes | 0 | 0 | 0 | 0 | 0 | 0.55 | 0 | 0 | 0 | 2.31 |
| 130 | P1093 | N#CCCNC(=O)Nc1ccc(cc1)Oc1ccnc(c1)C(=O)c1oc(c(c1)C)C | 404.42 | 0.46 | -9.483 | -8.184 | VAL 470, ALA 480, LYS 482, GLU 500, LEU 513, THR 528, PHE 582, PHE 594 | LYS 482, GLU 500, CYS 531, ASP 593 | - | TRP 530, PHE 594 | - | Yes | 0 | 0 | 0 | 0 | 0 | 0.55 | 0 | 0 | 2 | 3.4 |
| 131 | P1100 | O=C(Nc1ccc(cc1)Oc1ccnc(c1)C(=O)O)NC(=O)OC(Cl)C | 379.75 | 0.46 | -9.668 | -7.292 | LYS 482, GLU 500, VAL 503, THR 528, TRP 530 | LYS 482, GLU 500, CYS 531, ASP 593 | - | PHE 594 | - | Yes | 0 | 0 | 0 | 0 | 0 | 0.56 | 0 | 1 | 2 | 3.22 |
| 132 | P1110 | N#CC1(CC1)c1ccc(cc1)NC(=O)Nc1ccc(cc1)Oc1cncc(c1Cl)F | 422.84 | 0.41 | -10.473 | -7.851 | ALA 480, LYS 482, GLU 500, VAL 503, LEU 504, THR 528, LEU 566, ASP 593, PHE 594 | LYS 482, GLU 500, CYS 531, ASP 593 | - | TRP 530, PHE 594 | - | Yes | 0 | 1 | 0 | 0 | 0 | 0.55 | 0 | 0 | 2 | 2.89 |
| 133 | P1113 | N#Cc1ncc(cc1Cl)Oc1ccc(cc1)NC(=O)Nc1ccc(cc1)Nc1cnccc1C | 470.91 | 0.45 | -10.857 | -8.132 | ILE 462, VAL 470, LYS 482, GLU 500, LEU 504, LEU 513, THR 528, ILE 571, ASP 593 | LYS 482, GLU 500, CYS 531, ASP 593 | - | PHE 594 | - | Yes | 0 | 2 | 0 | 1 | 0 | 0.55 | 0 | 0 | 3 | 3.62 |
| 134 | P1114 | COc1ccc(cc1C(F)(F)F)NC(=O)Nc1ccc(cc1)Oc1cnccc1C | 417.38 | 0.57 | -11.381 | -8.045 | ILE 462, VAL 470, LYS 482, GLU 500, VAL 503, LEU 504, THR 528, ASP 593 | LYS 482, GLU 500, CYS 531, ASP 593 | - | PHE 594 | - | Yes | 0 | 1 | 0 | 1 | 0 | 0.55 | 0 | 0 | 3 | 3.01 |
| 135 | P1117 | O=C(Nc1ccc(cc1)Oc1ccnc(c1)C(=O)N(C)C)NC(=O)OC(=C)c1ccccc1 | 446.46 | 0.44 | -10.998 | -8.347 | ALA 480, LYS 482, GLU 500, VAL 503, LEU 504, THR 528, LEU 566, ILE 571 | LYS 482, GLU 500, CYS 531, ASP 593 | - | PHE 594 | - | Yes | 0 | 0 | 1 | 0 | 0 | 0.55 | 0 | 1 | 3 | 3.37 |
| 136 | P1131 | N#Cc1nccc(c1)Oc1ccc(cc1)NC(=O)NC(=O)C(F)(F)F | 350.25 | 0.5 | -9.347 | -8.099 | ALA 480, LYS 482, LEU 513, THR 528 | LYS 482, GLU 500, CYS 531, ASP 593 | - | PHE 594 | - | Yes | 0 | 0 | 0 | 0 | 0 | 0.55 | 0 | 0 | 1 | 2.38 |
| 137 | P1135 | O=Cc1nccc(c1)Oc1ccc(cc1)NC(=O)NCC(=O)Nc1ccc(cc1)C(N(C)C)C | 461.51 | 0.46 | -10.393 | -8.08 | ALA 480, LYS 482, VAL 503, LEU 513, THR 528, ILE 571 | LYS 482, GLU 500, CYS 531, ASP 593 | - | PHE 594 | - | Yes | 0 | 0 | 1 | 0 | 0 | 0.55 | 0 | 1 | 2 | 3.66 |

Supplementary Table 1: Molecule descriptors, Tanimoto similarity value, docking scores, target-ligand interactions, and ADME properties of 214 PURE ligands

| S. No. | Ligand | Canonical SMILES | Molecular weight (Da) | Tanimoto Similarity | Docking score with BRAF (kcal/mol) | Docking score with ABCG2 (kcal/mol) | Hydrophobic interactions | Hydrogen bonds | Salt bridge | Pi stack interaction(s) | Pi-cation interaction | Hydrogen bond match | Lipinski violation(s) | Ghose violation(s) | Veber violation(s) | Egan violation(s) | Muegge violation(s) | Bioavailability score | PAINS alert(s) | Brenk alert(s) | Leadlikeness violation(s) | Synthetic accessibility score |
|---|---|---|---|---|---|---|---|---|---|---|---|---|---|---|---|---|---|---|---|---|---|---|
| 138 | P1143 | COC(=O)c1cncc(c1)Oc1cc(ccc1N(C)C)NC(=O)Nc1ccc(cc1)Oc1ccncc1 | 499.52 | 0.43 | -10.429 | -8.274 | ALA 480, LYS 482, GLU 500, LEU 504, THR 528, PHE 594 | LYS 482, GLU 500, CYS 531, ASP 593 | - | PHE 594 | - | Yes | 0 | 2 | 1 | 0 | 0 | 0.55 | 1 | 0 | 2 | 3.76 |
| 139 | P1156 | COC(=O)c1nccc(c1)Oc1ccc(cc1)NC(=O)NC(=O)OC(=O)C | 373.32 | 0.44 | -9.406 | -6.958 | ALA 480, LYS 482, LEU 513, THR 528, ASP 593 | LYS 482, GLU 500, CYS 531, ASP 593 | - | PHE 594 | - | Yes | 0 | 0 | 0 | 1 | 0 | 0.55 | 0 | 2 | 2 | 2.85 |
| 140 | P1157 | O=C(NC(=O)C1CC1)Nc1ccc(cc1)Oc1ccnc(c1)C(=O)O | 341.32 | 0.45 | -9.916 | -8.106 | LYS 482, GLU 500, VAL 503, LEU 513, THR 528 | LYS 482, GLU 500, CYS 531, ASP 593 | - | PHE 594 | - | Yes | 0 | 0 | 0 | 0 | 0 | 0.56 | 0 | 0 | 1 | 2.43 |
| 141 | P1160 | CCOc1ccc(cc1Br)NC(=O)Nc1ccc(cc1)Oc1cccnc1 | 428.28 | 0.46 | -9.738 | -7.559 | GLU 500, LEU 504, THR 528, ILE 571, ASP 593, PHE 594 | LYS 482, GLU 500, CYS 531, ASP 593 | - | TRP 530, PHE 594 | - | Yes | 0 | 0 | 0 | 0 | 0 | 0.55 | 0 | 0 | 3 | 3.05 |
| 142 | P1164 | N=CCCNC(=O)Nc1ccccc1CNC(=O)c1nccc(c1)Oc1ccc(cc1)N(C(=O)C)O | 490.51 | 0.4 | -9.495 | -8.403 | ALA 480, LEU 504, LEU 513, THR 528 | LYS 482, GLU 500, CYS 531, GLY 533, ASP 593 | - | PHE 594 | - | Yes | 1 | 2 | 2 | 1 | 1 | 0.55 | 0 | 3 | 2 | 3.52 |
| 143 | P1173 | CNC(=O)Nc1ccc(cc1)Oc1ccnc(c1)C(=O)OC | 301.3 | 0.53 | -8.856 | -7.404 | ALA 480, LYS 482, LEU 513, THR 528 | LYS 482, GLU 500, CYS 531, ASP 593 | - | PHE 594 | - | Yes | 0 | 0 | 0 | 0 | 0 | 0.55 | 0 | 0 | 0 | 2.47 |
| 144 | P1181 | N#Cc1nccc(c1)Oc1ccc(cc1)NC(=O)NC(=O)OC(=O)C | 340.29 | 0.42 | -8.559 | -7.744 | VAL 470, VAL 503, LEU 504, TRP 530, PHE 594 | LYS 482, GLU 500, CYS 531, ASP 593 | - | PHE 594 | - | Yes | 0 | 0 | 0 | 0 | 0 | 0.55 | 0 | 2 | 1 | 2.65 |
| 145 | P1184 | O=C(Nc1ccc(cc1)C1(CC1)C(=O)O)Nc1ccc(cc1)Oc1cncc(c1Cl)F | 441.84 | 0.44 | -10.991 | -7.898 | GLU 500, VAL 503, LEU 504, LEU 513, THR 528, LEU 566, ILE 571, ASP 593 | LYS 482, GLU 500, CYS 531, HIS 573, ASP 593 | HIS 573 | TRP 530, PHE 594 | - | Yes | 0 | 0 | 0 | 0 | 0 | 0.56 | 0 | 0 | 2 | 2.94 |
| 146 | P1189 | ONC(=N)c1nccc(c1)Oc1ccc(cc1)NC(=O)NCC1CC1 | 341.36 | 0.43 | -9.875 | -8.37 | ALA 480, LYS 482, GLU 500, VAL 503, LEU 504, LEU 513, THR 528 | LYS 482, GLU 500, CYS 531, GLY 533, ASP 593 | - | PHE 594 | - | Yes | 0 | 0 | 0 | 0 | 0 | 0.55 | 0 | 3 | 1 | 2.73 |
| 147 | P1197 | O=NN(CCCN(c1ccc(cc1C)NC(=O)Nc1ccc(cc1)Oc1cncc(c1)Cl)C)C | 482.96 | 0.43 | -9.904 | -7.648 | LYS 482, GLU 500, VAL 503, LEU 504, LEU 513, THR 528 | LYS 482, GLU 500, CYS 531, ASP 593 | - | PHE 594 | - | Yes | 0 | 2 | 1 | 0 | 0 | 0.55 | 1 | 1 | 3 | 3.47 |
| 148 | P1206 | N#CC(c1ccc(cc1)NC(=O)Nc1ccc(cc1)Oc1cncc(c1NC)F)(C#N)C | 430.43 | 0.42 | -10.111 | -7.6 | LYS 482, GLU 500, VAL 503, LEU 504, THR 507, LEU 566, ASP 593, PHE 594 | LYS 482, GLU 500, CYS 531, ASP 593 | - | TRP 530, PHE 594 | - | Yes | 0 | 0 | 0 | 0 | 0 | 0.55 | 0 | 0 | 2 | 3.19 |
| 149 | P1207 | C#CCONC(=N)C(c1ccc(cc1)NC(=O)Nc1ccc(cc1)Oc1cccnc1)(C(=O)N)C | 472.5 | 0.41 | -10.127 | -8.339 | VAL 470, ALA 480, GLU 500, VAL 503, LEU 504, THR 528, LEU 566, PHE 582, ASP 593, PHE 594 | LYS 482, GLU 500, VAL 503, CYS 531, ASP 593 | - | TRP 530, PHE 594 | - | Yes | 0 | 1 | 2 | 1 | 1 | 0.55 | 0 | 4 | 2 | 4.2 |
| 150 | P1228 | N#Cc1nccc(c1)Oc1ccc(cc1)NC(=O)NC(=O)OC(Cl)C | 360.75 | 0.43 | -9.285 | -7.193 | ALA 480, LYS 482, VAL 503, THR 528 | LYS 482, GLU 500, CYS 531, ASP 593 | - | PHE 594 | - | Yes | 0 | 0 | 0 | 0 | 0 | 0.55 | 0 | 1 | 2 | 3.22 |
| 151 | P1237 | N#Cc1nccc(c1)Oc1ccc(cc1)NC(=O)NCC(=O)Nc1cccc(c1)Br | 466.29 | 0.5 | -10.547 | -8.215 | LYS 482, ASN 499, GLU 500, VAL 503, LEU 513, THR 528 | LYS 482, GLU 500, CYS 531, ASP 593 | - | PHE 594 | - | Yes | 0 | 0 | 0 | 0 | 0 | 0.55 | 0 | 0 | 2 | 3.05 |
| 152 | P1240 | CCOC(=O)NC(=O)Nc1ccc(cc1)Oc1ccnc(c1)C#N | 326.31 | 0.43 | -9.002 | -7.608 | ALA 480, LYS 482, VAL 503, LEU 513, THR 528 | LYS 482, GLU 500, CYS 531, ASP 593 | - | PHE 594 | - | Yes | 0 | 0 | 0 | 0 | 0 | 0.55 | 0 | 0 | 1 | 2.66 |
| 153 | P1262 | CNCCCN(c1ccc(cc1C(=O)C)NC(=O)Nc1ccc(cc1)Oc1cncc(c1)Cl)C=O | 495.96 | 0.47 | -9.848 | -7.718 | LYS 482, GLU 500, LEU 504, TRP 530, LEU 566, PHE 594 | LYS 482, GLU 500, LEU 513, CYS 531, ASP 593 | - | PHE 594 | - | Yes | 0 | 2 | 1 | 0 | 0 | 0.55 | 0 | 1 | 2 | 3.49 |
| 154 | P1266 | FCc1nccc(c1)Oc1ccc(cc1)NC(=O)NC(=O)OC(Br)C | 412.21 | 0.43 | -9.387 | -7.283 | ALA 480, LYS 482, GLU 500, VAL 503, LEU 504, LEU 513, THR 528 | LYS 482, GLU 500, CYS 531, ASP 593 | - | PHE 594 | - | Yes | 0 | 0 | 0 | 0 | 0 | 0.55 | 0 | 1 | 3 | 3.32 |
| 155 | P1277 | N=CCCNC(=O)Nc1ccc(cc1)Oc1ccnc(c1)C(=O)NCc1ccccc1 | 417.46 | 0.48 | -9.991 | -8.41 | ALA 480, LYS 482, LEU 513, THR 528, ASP 593 | LYS 482, GLU 500, CYS 531, GLY 592, ASP 593 | - | PHE 582, PHE 594 | - | Yes | 0 | 0 | 1 | 0 | 0 | 0.55 | 0 | 1 | 2 | 2.99 |
| 156 | P1279 | N#Cc1nccc(c1)Oc1ccc(cc1)NC(=O)NC(=O)Cc1ccc(c(c1)OC)Br | 481.3 | 0.46 | -9.746 | -7.906 | GLU 500, THR 528, TRP 530, ASP 593, PHE 594 | LYS 482, GLU 500, CYS 531, ASP 593 | - | PHE 594 | - | Yes | 0 | 1 | 0 | 0 | 0 | 0.55 | 0 | 0 | 3 | 2.95 |
| 157 | P1287 | N#CC(c1ccc(cc1)NC(=O)Nc1ccc(cc1)Oc1cncc(c1)Cl)(C#N)C | 417.85 | 0.45 | -10.165 | -7.726 | ALA 480, GLU 500, VAL 503, LEU 504, THR 528, TRP 530, LEU 566, ASP 593, PHE 594 | LYS 482, GLU 500, CYS 531, ASP 593 | - | PHE 594 | - | Yes | 0 | 0 | 0 | 0 | 0 | 0.55 | 0 | 0 | 2 | 2.87 |
| 158 | P1294 | CNc1ccc(cc1)NC(=O)CNC(=O)Nc1ccc(cc1)Oc1ccnc(c1)C#N | 416.43 | 0.5 | -10.043 | -8.121 | ALA 480, LYS 482, VAL 503, THR 528 | LYS 482, GLU 500, CYS 531, ILE 572, ASP 593 | - | PHE 594 | HIS 573 | Yes | 0 | 0 | 0 | 0 | 0 | 0.55 | 0 | 0 | 2 | 3.02 |
| 159 | P1297 | O=CN(c1ccc(cc1)NC(=O)Nc1ccc(cc1)Oc1cncc(c1CO)F)C | 410.4 | 0.41 | -10.117 | -8.076 | LYS 482, GLU 500, LEU 504, THR 528, ASP 593, PHE 594 | LYS 482, GLU 500, CYS 531, ASP 593 | - | TRP 530, PHE 594 | - | Yes | 0 | 0 | 0 | 0 | 0 | 0.55 | 0 | 1 | 2 | 3.04 |
| 160 | P1315 | CNc1ccc(cc1C(F)(F)F)NC(=O)Nc1ccc(cc1)Oc1cccnc1 | 402.37 | 0.63 | -11.087 | -7.839 | LYS 482, LEU 504, LEU 513, THR 528, TRP 530, ASP 593 | LYS 482, GLU 500, CYS 531, ASP 593 | - | PHE 594 | - | Yes | 0 | 1 | 0 | 1 | 0 | 0.55 | 0 | 0 | 3 | 2.9 |
| 161 | P1321 | O=C(Nc1ccc(c(c1)Cl)N(C)C)Nc1ccc(cc1)Oc1cccnc1 | 382.84 | 0.51 | -10.009 | -7.46 | LYS 482, GLU 500, LEU 504, THR 528 | LYS 482, GLU 500, CYS 531, ASP 593 | - | TRP 530, PHE 594 | - | Yes | 0 | 0 | 0 | 0 | 0 | 0.55 | 1 | 0 | 2 | 2.9 |
| 162 | P1327 | O=NN(c1ccc(cc1)NC(=O)Nc1ccc(cc1)Oc1cncc(c1)C(=O)OC)C | 421.41 | 0.44 | -9.951 | -7.735 | LYS 482, GLU 500, LEU 504, THR 528, TRP 530, ASP 593, PHE 594 | LYS 482, GLU 500, CYS 531, HIS 573, ASP 593 | - | PHE 594 | - | Yes | 0 | 0 | 0 | 0 | 0 | 0.55 | 0 | 1 | 2 | 3.21 |
| 163 | P1329 | COC(=O)CNC(=O)Nc1ccc(cc1)Oc1ccnc(c1)N(C)C | 344.37 | 0.42 | -8.88 | -7.303 | ALA 480, LYS 482, LEU 513, THR 528 | LYS 482, GLU 500, CYS 531, ASP 593 | - | PHE 594 | - | Yes | 0 | 0 | 0 | 0 | 0 | 0.55 | 0 | 0 | 1 | 2.82 |
| 164 | P1341 | NCc1ccccc1S(=O)(=O)OC(=O)NC(=O)Nc1ccc(cc1)Oc1ccnc(c1)CCl | 490.92 | 0.42 | -10.487 | -8.276 | ALA 480, LYS 482, VAL 503, LEU 513, THR 528, LEU 566, ILE 571 | LYS 482, GLU 500, CYS 531, ASP 593 | - | PHE 594 | HIS 573 | Yes | 0 | 1 | 2 | 1 | 1 | 0.55 | 0 | 2 | 2 | 3.46 |
| 165 | P1362 | O=C(Nc1ccc(cc1)Oc1ccnc(c1)N)NC(=O)OC(Br)C | 395.21 | 0.43 | -9.047 | -7.361 | ALA 480, LYS 482, GLU 500, VAL 503, THR 528, TRP 530 | LYS 482, GLU 500, CYS 531, ASP 593 | - | PHE 594 | - | Yes | 0 | 0 | 0 | 0 | 0 | 0.55 | 0 | 1 | 2 | 3.41 |

Supplementary Table 1: Molecule descriptors, Tanimoto similarity value, docking scores, target-ligand interactions, and ADME properties of 214 PURE ligands

| S. No. | Ligand | Canonical SMILES | Molecular weight (Da) | Tanimoto Similarity | Docking score with BRAF (kcal/mol) | Docking score with ABCG2 (kcal/mol) | Hydrophobic interactions | Hydrogen bonds | Salt bridge | Pi stack interaction(s) | Pi-cation interaction | Hydrogen bond match | Lipinski violation(s) | Ghose violation(s) | Veber violation(s) | Egan violation(s) | Muegge violation(s) | Bioavailability score | PAINS alert(s) | Brenk alert(s) | Leadlikeness violation(s) | Synthetic accessibility score |
|---|---|---|---|---|---|---|---|---|---|---|---|---|---|---|---|---|---|---|---|---|---|---|
| 166 | P1367 | CCOC(=O)NC(=O)Nc1ccc(cc1)Oc1ccnc(c1)N(CC)C | 358.39 | 0.41 | -8.816 | -7.422 | ALA 480, GLU 500, VAL 503, THR 528, PHE 582 | LYS 482, GLU 500, CYS 531, ASP 593 | - | TRP 530, PHE 594 | - | Yes | 0 | 0 | 0 | 0 | 0 | 0.55 | 0 | 0 | 2 | 3.07 |
| 167 | P1371 | CNCCCN(c1ccc(cc1CN)NC(=O)Nc1ccc(cc1)Oc1cncc(c1)O)C | 450.53 | 0.43 | -9.206 | -7.53 | LYS 482, GLU 500, LEU 504, THR 528 | LYS 482, GLU 500, LEU 513, CYS 531, HIS 573, ILE 591, ASP 593 | - | PHE 594 | - | Yes | 0 | 1 | 1 | 0 | 0 | 0.55 | 1 | 0 | 2 | 3.48 |
| 168 | P1372 | NN(c1ccc(cc1)Cl)CCCC(=O)NCc1nccc(c1)Oc1ccc(cc1)NC(=O)N | 468.94 | 0.41 | -9.383 | -8.151 | ILE 462, VAL 470, LYS 482, LEU 513, TRP 530, PHE 582, PHE 594, LEU 596 | ILE 462, LYS 482, GLU 500, CYS 531, ASP 593 | - | PHE 594 | - | Yes | 0 | 0 | 1 | 1 | 0 | 0.55 | 0 | 1 | 2 | 3.08 |
| 169 | P1379 | O=CN(c1ccc(cc1)NC(=O)Nc1ccc(cc1)Oc1cncc(c1)C(O)C)C | 406.43 | 0.43 | -10.702 | -7.725 | ALA 480, LYS 482, GLU 500, LEU 504, THR 528, TRP 530, PHE 582 | LYS 482, GLU 500, CYS 531, ASP 593 | - | PHE 594 | - | Yes | 0 | 0 | 0 | 0 | 0 | 0.55 | 0 | 1 | 2 | 3.54 |
| 170 | P1384 | O=CNNc1ccc(cc1)Oc1ccnc(c1)CNC(=O)c1ccccc1Cl | 396.83 | 0.44 | -9.445 | -8.318 | ALA 480, LYS 482, LEU 513, THR 528, PHE 582 | LYS 482, GLU 500, CYS 531, ASP 593 | - | PHE 594 | - | Yes | 0 | 0 | 0 | 0 | 0 | 0.55 | 0 | 1 | 2 | 2.71 |
| 171 | P1389 | CNC(=O)c1ccc(o1)CNc1ccc(cc1)Oc1ccnc(c1)CN | 352.39 | 0.44 | -9.415 | -7.901 | ALA 480, LYS 482, GLU 500, THR 528, ASP 593 | LYS 482, GLU 500, CYS 531, ILE 591, ASP 593 | - | PHE 594 | - | Yes | 0 | 0 | 0 | 0 | 0 | 0.55 | 0 | 0 | 2 | 2.97 |
| 172 | P1394 | O=C(Nc1ccc(cc1)F)Nc1ccc(cc1)Oc1cccnc1 | 323.32 | 0.46 | -10.246 | -7.835 | ALA 480, LYS 482, GLU 500, VAL 503, LEU 504, THR 528 | LYS 482, GLU 500, CYS 531, ASP 593 | - | PHE 594 | - | Yes | 0 | 0 | 0 | 0 | 0 | 0.55 | 0 | 0 | 0 | 2.54 |
| 173 | P1396 | CNC(=O)c1nccc(c1)Oc1cc(ccc1F)NC(=O)NC(=O)OC(=O)C | 390.32 | 0.44 | -9.558 | -8.11 | VAL 470, ALA 480, LYS 482, GLU 500, VAL 503, LEU 513, ILE 526, THR 528, PHE 594 | LYS 482, GLU 500, CYS 531, ASP 593 | - | - | - | Yes | 0 | 0 | 0 | 1 | 0 | 0.55 | 0 | 2 | 2 | 2.89 |
| 174 | P1402 | O=C(NC(=O)c1ccccc1)Nc1ccc(cc1)Oc1ccnc(c1)C(N)C | 376.41 | 0.47 | -10.954 | -8.371 | VAL 470, LYS 482, GLU 500, VAL 503, LEU 504, LEU 513, TRP 530, PHE 582, ASP 593 | LYS 482, GLU 500, CYS 531, ASP 593 | - | PHE 594 | - | Yes | 0 | 0 | 0 | 0 | 0 | 0.55 | 0 | 0 | 2 | 3.11 |
| 175 | P1412 | NCC(c1ccc(cc1)NC(=O)Nc1ccc(cc1)Oc1cncc(c1)S)C=O | 408.47 | 0.42 | -9.43 | -7.159 | LYS 482, GLU 500, VAL 503, LEU 504, THR 528, TRP 530, ASP 593, PHE 594 | LYS 482, GLU 500, THR 507, CYS 531, ASP 593 | - | PHE 594 | - | Yes | 0 | 0 | 1 | 1 | 0 | 0.55 | 0 | 2 | 2 | 3.4 |
| 176 | P1434 | CNCCCN(c1ccc(cc1C)NC(=O)Nc1ccc(cc1)Oc1cncc(c1)O)C | 435.52 | 0.44 | -9.717 | -7.568 | LYS 482, GLU 500, VAL 503, LEU 504, TRP 530, ILE 571, ASP 593, PHE 594 | LYS 482, GLU 500, CYS 531, HIS 573, ASP 593 | - | PHE 594 | - | Yes | 0 | 0 | 1 | 0 | 0 | 0.55 | 1 | 0 | 2 | 3.34 |
| 177 | P1445 | COC(=O)c1nccc(c1)Oc1ccc(cc1)NC(=O)NC(=O)C(=CN(C)C)c1ccccc1 | 460.48 | 0.43 | -10.657 | -8.037 | ALA 480, LYS 482, VAL 503, LEU 504, THR 528, LEU 566, ASP 593, PHE 594 | LYS 482, GLU 500, CYS 531, ASP 593 | - | PHE 594 | - | Yes | 0 | 0 | 1 | 0 | 0 | 0.55 | 0 | 1 | 3 | 3.5 |
| 178 | P1449 | COC(=O)c1cncc(c1)Oc1ccc(cc1)NC(=O)Nc1ccc(cc1)F | 381.36 | 0.48 | -10.539 | -7.691 | LYS 482, GLU 500, VAL 503, LEU 504, LEU 513, THR 528 | LYS 482, GLU 500, CYS 531, ASP 593 | - | PHE 594 | - | Yes | 0 | 0 | 0 | 0 | 0 | 0.55 | 0 | 0 | 2 | 2.84 |
| 179 | P1465 | O=C(Nc1ccc(c(c1)c1ccnc(c1)C)N(C)C)Nc1ccc(cc1)Oc1cnccc1C | 453.54 | 0.42 | -11.543 | -8.378 | ILE 462, VAL 470, ASN 499, GLU 500, VAL 503, LEU 504, THR 528, ASP 593 | LYS 482, GLU 500, CYS 531, ASP 593 | - | TRP 530, PHE 594 | - | Yes | 0 | 2 | 0 | 1 | 0 | 0.55 | 1 | 0 | 3 | 3.53 |
| 180 | P1467 | CCOC(=O)COc1ccc(cc1C)OC(c1nccc(c1)Oc1ccc(cc1)NC(=O)NC)C | 479.52 | 0.45 | -9.332 | -8.009 | ILE 462, VAL 470, ALA 480, LYS 482, LEU 513, THR 528, TRP 530, PHE 582, PHE 594 | LYS 482, GLU 500, CYS 531, ASP 593 | - | PHE 594 | - | Yes | 0 | 1 | 1 | 0 | 0 | 0.55 | 0 | 0 | 3 | 4.2 |
| 181 | P1479 | N#Cc1nccc(c1)Oc1ccc(cc1)NC(=O)NC(=O)Cc1ccc(c(c1)OC)N | 417.42 | 0.46 | -10.065 | -7.781 | GLU 500, THR 528, ASP 593, PHE 594 | LYS 482, GLU 500, CYS 531, ASP 593 | - | TRP 530, PHE 594 | - | Yes | 0 | 0 | 0 | 1 | 0 | 0.55 | 0 | 1 | 2 | 2.99 |
| 182 | P1507 | COC(=O)c1nccc(c1)Oc1ccc(cc1)NC(=O)NC(=O)c1c(Cl)cccc1Cl | 460.27 | 0.54 | -10.583 | -8.201 | ALA 480, LYS 482, VAL 503, LEU 513, THR 528, ASP 593 | LYS 482, GLU 500, CYS 531, ASP 593 | - | PHE 594 | - | Yes | 0 | 0 | 0 | 0 | 0 | 0.55 | 0 | 0 | 3 | 2.88 |
| 183 | P1517 | O=C(Nc1ccc(cc1)Cl)Nc1ccc(cc1)Oc1cncc(c1)Cl | 374.22 | 0.48 | -10.346 | -7.723 | LYS 482, GLU 500, VAL 503, LEU 504, THR 528 | LYS 482, GLU 500, CYS 531, ASP 593 | - | PHE 594 | - | Yes | 0 | 0 | 0 | 0 | 0 | 0.55 | 0 | 0 | 2 | 2.71 |
| 184 | P1523 | C=COC(=O)NC(=O)Nc1ccc(cc1)Oc1ccnc(c1)C(=O)OC | 357.32 | 0.45 | -8.993 | -7.481 | VAL 470, LYS 482, GLU 500, VAL 503, LEU 513, THR 528 | LYS 482, GLU 500, CYS 531, ASP 593 | - | PHE 594 | - | Yes | 0 | 0 | 0 | 0 | 0 | 0.55 | 0 | 2 | 2 | 2.85 |
| 185 | P1528 | N#CCCNC(=O)Nc1ccc(cc1)Oc1ccnc(c1)C(=O)OC | 340.33 | 0.45 | -8.627 | -6.829 | ALA 480, LYS 482, GLU 500, LEU 513, THR 528 | LYS 482, GLU 500, CYS 531, ASP 593 | - | PHE 594 | - | Yes | 0 | 0 | 0 | 0 | 0 | 0.55 | 0 | 0 | 1 | 2.66 |
| 186 | P1532 | O=C(Nc1ccc(cc1)Oc1ccnc(c1)C(=O)C)NC(=O)OC(Br)C | 422.23 | 0.45 | -9.549 | -7.401 | ALA 480, GLU 500, LEU 513, THR 528, TRP 530, ASP 593 | LYS 482, GLU 500, CYS 531, ASP 593 | - | PHE 594 | - | Yes | 0 | 0 | 0 | 0 | 0 | 0.55 | 0 | 1 | 2 | 3.34 |
| 187 | P1539 | N#CCCNC(=O)Nc1ccc(cc1)Oc1ccnc(c1)C(=O)N | 325.32 | 0.45 | -8.969 | -7.377 | VAL 470, ALA 480, LYS 482, GLU 500, LEU 513, THR 528 | LYS 482, GLU 500, CYS 531, ASP 593 | - | PHE 594 | - | Yes | 0 | 0 | 0 | 0 | 0 | 0.55 | 0 | 0 | 1 | 2.42 |
| 188 | P1564 | O=C(Nc1ccc(c(c1)C(F)(F)F)N(C)C)Nc1ccc(cc1)Oc1ccncc1 | 416.4 | 0.6 | -10.518 | -7.797 | ALA 480, LYS 482, LEU 504, LEU 513, THR 528, ASP 593 | LYS 482, GLU 500, CYS 531, ASP 593 | - | PHE 594 | - | Yes | 0 | 1 | 0 | 1 | 0 | 0.55 | 1 | 0 | 3 | 2.81 |
| 189 | P1573 | CCNC(=O)Nc1ccc(cc1)Oc1ccnc(c1)C#N | 282.3 | 0.42 | -8.689 | -7.073 | ALA 480, LYS 482, GLU 500, LEU 504, THR 528 | LYS 482, GLU 500, CYS 531, ASP 593 | - | PHE 594 | - | Yes | 0 | 0 | 0 | 0 | 0 | 0.55 | 0 | 0 | 0 | 2.37 |
| 190 | P1583 | C=COC(=O)NC(=O)Nc1ccc(cc1)Oc1ccnc(c1)CN | 328.32 | 0.41 | -8.949 | -7.649 | ALA 480, LYS 482, GLU 500, VAL 503, LEU 513, THR 528, PHE 594 | LYS 482, GLU 500, CYS 531, ASP 593 | - | PHE 594 | - | Yes | 0 | 0 | 0 | 0 | 0 | 0.55 | 0 | 1 | 1 | 2.66 |
| 191 | P1585 | O=C(Nc1ccc(cc1)C(C(=O)O)(C)C)Nc1ccc(cc1)Oc1cnccc1C | 405.45 | 0.44 | -10.369 | -8.352 | ILE 462, VAL 470, ALA 480, GLU 500, VAL 503, LEU 504, THR 528, LEU 566, PHE 582, ASP 593, PHE 594 | LYS 482, GLU 500, CYS 531, HIS 573, ASP 593 | HIS 573 | TRP 530, PHE 594 | - | Yes | 0 | 0 | 0 | 0 | 0 | 0.56 | 0 | 0 | 3 | 3.01 |
| 192 | P1594 | N=CCCNC(=O)Nc1ccc(cc1)Oc1ccnc(c1)C(=O)Cc1cccc(c1)Cl | 436.89 | 0.5 | -10.427 | -8.366 | ILE 462, ALA 480, LYS 482, LEU 513, THR 528, ASP 593, PHE 594 | LYS 482, GLU 500, CYS 531, GLY 592, ASP 593 | - | PHE 582, PHE 594 | - | Yes | 0 | 0 | 1 | 0 | 0 | 0.55 | 0 | 1 | 2 | 3.13 |
| 193 | P1601 | CCC(=O)NC1(CC1)c1ccc(cc1)NC(=O)Nc1ccc(cc1)Oc1cccnc1 | 416.47 | 0.44 | -10.517 | -7.738 | ALA 480, GLU 500, VAL 503, LEU 504, THR 528, LEU 566, PHE 582, ASP 593, PHE 594 | LYS 482, GLU 500, CYS 531, ASP 593 | - | TRP 530, PHE 594 | - | Yes | 0 | 0 | 0 | 0 | 0 | 0.55 | 0 | 0 | 2 | 2.87 |
| 194 | P1610 | O=C(NC(=O)c1ccccc1O)Nc1ccc(cc1)Oc1ccnc(c1)C(N)C | 392.41 | 0.47 | -10.862 | -8.118 | VAL 470, ALA 480, LYS 482, GLU 500, VAL 503, LEU 513, THR 528, TRP 530, PHE 582, ASP 593 | LYS 482, GLU 500, CYS 531, ASP 593 | - | PHE 594 | - | Yes | 0 | 0 | 0 | 0 | 0 | 0.55 | 0 | 0 | 2 | 3.24 |

### Supplementary Table 1: Molecule descriptors, Tanimoto similarity value, docking scores, target-ligand interactions, and ADME properties of 214 PURE ligands

| S. No. | Ligand | Canonical SMILES | Molecular weight (Da) | Tanimoto Similarity | Docking score with BRAF (kcal/mol) | Docking score with ABCG2 (kcal/mol) | Hydrophobic interactions | Hydrogen bonds | Salt bridge | Pi stack interaction(s) | Pi-cation interaction | Hydrogen bond match | Lipinski violation(s) | Ghose violation(s) | Veber violation(s) | Egan violation(s) | Muegge violation(s) | Bioavailability score | PAINS alert(s) | Brenk alert(s) | Leadlikeness violation(s) | Synthetic accessibility score |
|---|---|---|---|---|---|---|---|---|---|---|---|---|---|---|---|---|---|---|---|---|---|---|
| 195 | P1626 | COC(=O)c1cncc(c1)Oc1ccc(cc1)NC(=O)Nc1ccc(cc1)N(C)C | 406.43 | 0.45 | -10.101 | -7.878 | GLU 500, LEU 504, THR 528, ASP 593 | LYS 482, GLU 500, CYS 531, ASP 593 | - | PHE 594 | - | Yes | 0 | 0 | 0 | 0 | 0 | 0.55 | 1 | 0 | 2 | 3.15 |
| 196 | P1628 | NCC(c1ccc(cc1)NC(=O)Nc1ccc(cc1)Oc1cncc(c1)F)C=O | 394.4 | 0.43 | -9.681 | -7.286 | LYS 482, GLU 500, VAL 503, LEU 504, THR 528, PHE 594 | LYS 482, GLU 500, THR 507, CYS 531, ASP 593 | - | PHE 594 | - | Yes | 0 | 0 | 0 | 0 | 0 | 0.55 | 0 | 1 | 2 | 3.32 |
| 197 | P1641 | O=C(Nc1ccc(cc1)Oc1ccnc(c1)C(=O)N(C)C)NCC(=O)C | 356.38 | 0.45 | -9.507 | -7.763 | ALA 480, LYS 482, GLU 500, VAL 503, LEU 504, THR 528 | LYS 482, GLU 500, CYS 531, ASP 593 | - | PHE 594 | - | Yes | 0 | 0 | 0 | 0 | 0 | 0.55 | 0 | 0 | 2 | 2.6 |
| 198 | P1651 | CCOc1ccc(cc1)NC(=O)CNC(=O)Nc1ccc(cc1)Oc1ccnc(c1)C=O | 434.44 | 0.47 | -9.993 | -7.837 | ALA 480, LYS 482, VAL 503, LEU 513, THR 528, ILE 571 | LYS 482, GLU 500, CYS 531, ASP 593 | - | PHE 594 | - | Yes | 0 | 0 | 1 | 0 | 0 | 0.55 | 0 | 1 | 2 | 2.99 |
| 199 | P1662 | CNCCCN(c1ccc(cc1C(=O)NO)NC(=O)Nc1ccc(cc1)Oc1cncc(c1)Cl)C | 498.96 | 0.48 | -9.433 | -8.11 | LYS 482, GLU 500, VAL 503, LEU 504, LEU 513, THR 528, ASP 593 | LYS 482, GLU 500, CYS 531, ASP 593 | - | PHE 594 | - | Yes | 0 | 2 | 1 | 0 | 0 | 0.55 | 1 | 2 | 2 | 3.61 |
| 200 | P1675 | COC(=O)c1cncc(c1)Oc1ccc(cc1)NC(=O)Nc1ccc(cc1)Br | 442.26 | 0.47 | -10.163 | -7.81 | LYS 482, GLU 500, VAL 503, LEU 504, LEU 513, THR 528 | LYS 482, GLU 500, CYS 531, ASP 593 | - | PHE 594 | - | Yes | 0 | 0 | 0 | 0 | 0 | 0.55 | 0 | 0 | 2 | 2.92 |
| 201 | P1682 | C=CCNC(=O)Nc1ccc(cc1)Oc1ccnc(c1)C(=O)C(C)C | 339.39 | 0.46 | -9.242 | -7.457 | ILE 462, LYS 482, GLU 500, VAL 503, LEU 504, THR 528, PHE 582, PHE 594 | LYS 482, GLU 500, CYS 531, ASP 593 | - | PHE 594 | - | Yes | 0 | 0 | 0 | 0 | 0 | 0.55 | 0 | 1 | 1 | 2.75 |
| 202 | P1683 | O=C(Nc1ccc(cc1)Oc1ccncc1)Nc1ccc(c(c1)c1ccc(cn1)CC(=O)O)N(C)C | 483.52 | 0.44 | -10.421 | -8.099 | GLU 500, VAL 503, LEU 504, THR 528, ASP 593, PHE 594 | LYS 482, GLU 500, CYS 531, HIS 573, ASP 593 | ARG 574 | TRP 530, PHE 594 | - | Yes | 0 | 2 | 0 | 0 | 0 | 0.56 | 1 | 0 | 2 | 3.45 |
| 203 | P1686 | CNc1ccc(cc1Br)NC(=O)Nc1ccc(cc1)Oc1cccnc1 | 413.27 | 0.5 | -10.244 | -7.633 | LYS 482, GLU 500, VAL 503, LEU 504, LEU 513, THR 528, ASP 593 | LYS 482, GLU 500, CYS 531, ASP 593 | - | PHE 594 | - | Yes | 0 | 0 | 0 | 0 | 0 | 0.55 | 0 | 0 | 2 | 3 |
| 204 | P1688 | O=C(Nc1ccc(c(c1)C(=O)O)N(C)C)Nc1ccc(cc1)Oc1cccnc1 | 392.41 | 0.49 | -10.043 | -7.603 | LYS 482, GLU 500, VAL 503, LEU 504, THR 528 | LYS 482, GLU 500, LEU 513, CYS 531, ASP 593 | - | PHE 594 | - | Yes | 0 | 0 | 0 | 0 | 0 | 0.56 | 1 | 0 | 2 | 2.89 |
| 205 | P1689 | CNC(=O)Nc1ccc(cc1)Oc1ccnc(c1)C#N | 268.27 | 0.46 | -8.637 | -6.996 | LYS 482, LEU 513, THR 528 | LYS 482, GLU 500, CYS 531, ASP 593 | - | PHE 594 | - | Yes | 0 | 0 | 0 | 0 | 0 | 0.55 | 0 | 0 | 0 | 2.27 |
| 206 | P1690 | N#CC1(CC1)c1ccc(cc1)NC(=O)Nc1ccc(cc1)Oc1cncc(c1)Cl | 404.85 | 0.43 | -10.46 | -7.628 | ALA 480, GLU 500, VAL 503, LEU 504, LEU 513, THR 528, TRP 530, LEU 566, ILE 571, ASP 593 | LYS 482, GLU 500, CYS 531, ASP 593 | - | PHE 594 | - | Yes | 0 | 0 | 0 | 0 | 0 | 0.55 | 0 | 0 | 2 | 2.86 |
| 207 | P1692 | N#Cc1nccc(c1)Oc1ccc(cc1)NC(=O)NC(=O)C(F)F | 332.26 | 0.44 | -9.478 | -8.027 | ALA 480, LYS 482, THR 528 | LYS 482, GLU 500, CYS 531, ASP 593 | - | PHE 594 | - | Yes | 0 | 0 | 0 | 0 | 0 | 0.55 | 0 | 0 | 0 | 2.38 |
| 208 | P1711 | OCCNC(=O)c1cncc(c1)Oc1ccc(cc1)NC(=O)Nc1ccc(cc1)N(C=O)C | 449.46 | 0.44 | -10.784 | -8.022 | LYS 482, GLU 500, LEU 504, THR 528, TRP 530, PHE 594 | LYS 482, GLU 500, CYS 531, GLY 533, ASP 593 | - | PHE 594 | - | Yes | 0 | 0 | 1 | 1 | 0 | 0.55 | 0 | 1 | 2 | 3.18 |
| 209 | P1718 | COC(=O)NC(=O)Nc1ccc(cc1)Oc1ccnc(c1)N(C)C | 330.34 | 0.43 | -9.289 | -7.414 | ALA 480, LYS 482, LEU 513, THR 528 | LYS 482, GLU 500, CYS 531, ASP 593 | - | PHE 594 | - | Yes | 0 | 0 | 0 | 0 | 0 | 0.55 | 0 | 0 | 1 | 2.8 |
| 210 | P1723 | O=C(NC(=O)c1ccccc1O)Nc1ccc(cc1)Oc1ccnc(c1)C(=O)C | 391.38 | 0.51 | -10.423 | -8.183 | ILE 462, LYS 482, GLU 500, VAL 503, LEU 504, THR 528, ASP 593 | LYS 482, GLU 500, CYS 531, HIS 573, ASP 593 | - | PHE 594 | - | Yes | 0 | 0 | 0 | 0 | 0 | 0.55 | 0 | 0 | 3 | 2.7 |
| 211 | P1727 | COC(=O)c1cncc(c1)Oc1cc(ccc1N(C)C)NC(=O)Nc1ccc(cc1)Oc1cccnc1 | 499.52 | 0.42 | -10.644 | -8.392 | LYS 482, GLU 500, LEU 504, LEU 513, THR 528, ASP 593 | LYS 482, GLU 500, CYS 531, ASP 593 | - | TRP 530, PHE 594 | - | Yes | 0 | 2 | 1 | 0 | 0 | 0.55 | 1 | 0 | 2 | 3.74 |
| 212 | P1737 | CCNC(=O)Nc1ccc(cc1)Oc1ccnc(c1)C(=O)N(C)C | 328.37 | 0.47 | -9.29 | -7.401 | ALA 480, LYS 482, GLU 500, THR 528 | LYS 482, GLU 500, CYS 531, ASP 593 | - | PHE 594 | - | Yes | 0 | 0 | 0 | 0 | 0 | 0.55 | 0 | 0 | 1 | 2.54 |
| 213 | P1743 | O=C(Nc1ccc(c(c1)Br)N(C)C)Nc1ccc(cc1)Oc1ccncc1 | 427.29 | 0.48 | -10.151 | -7.291 | ALA 480, LYS 482, GLU 500, LEU 504, THR 528, ASP 593 | LYS 482, GLU 500, CYS 531, ASP 593 | - | TRP 530, PHE 594 | - | Yes | 0 | 0 | 0 | 0 | 0 | 0.55 | 1 | 0 | 2 | 2.88 |
| 214 | P1749 | CN(Cc1ccncc1)Cc1ccc(cc1)NC(=O)Nc1ccc(cc1)Oc1cncc(c1)Cl | 473.95 | 0.41 | -10.178 | -8.217 | LYS 482, GLU 500, VAL 503, LEU 504, THR 528, TRP 530, PHE 594 | LYS 482, GLU 500, CYS 531, ASP 593 | - | PHE 594 | - | Yes | 0 | 1 | 0 | 0 | 0 | 0.55 | 0 | 0 | 3 | 3.29 |
