## Supplementart Figure 1 for "PURE: Policy-guided Unbiased REpresentations for structure-constrained molecular generation"

Supplementary Figure 1: Synthesis Pathways generated by PURE leading to the "top 9" molecules as alternatives to Sorafenib.

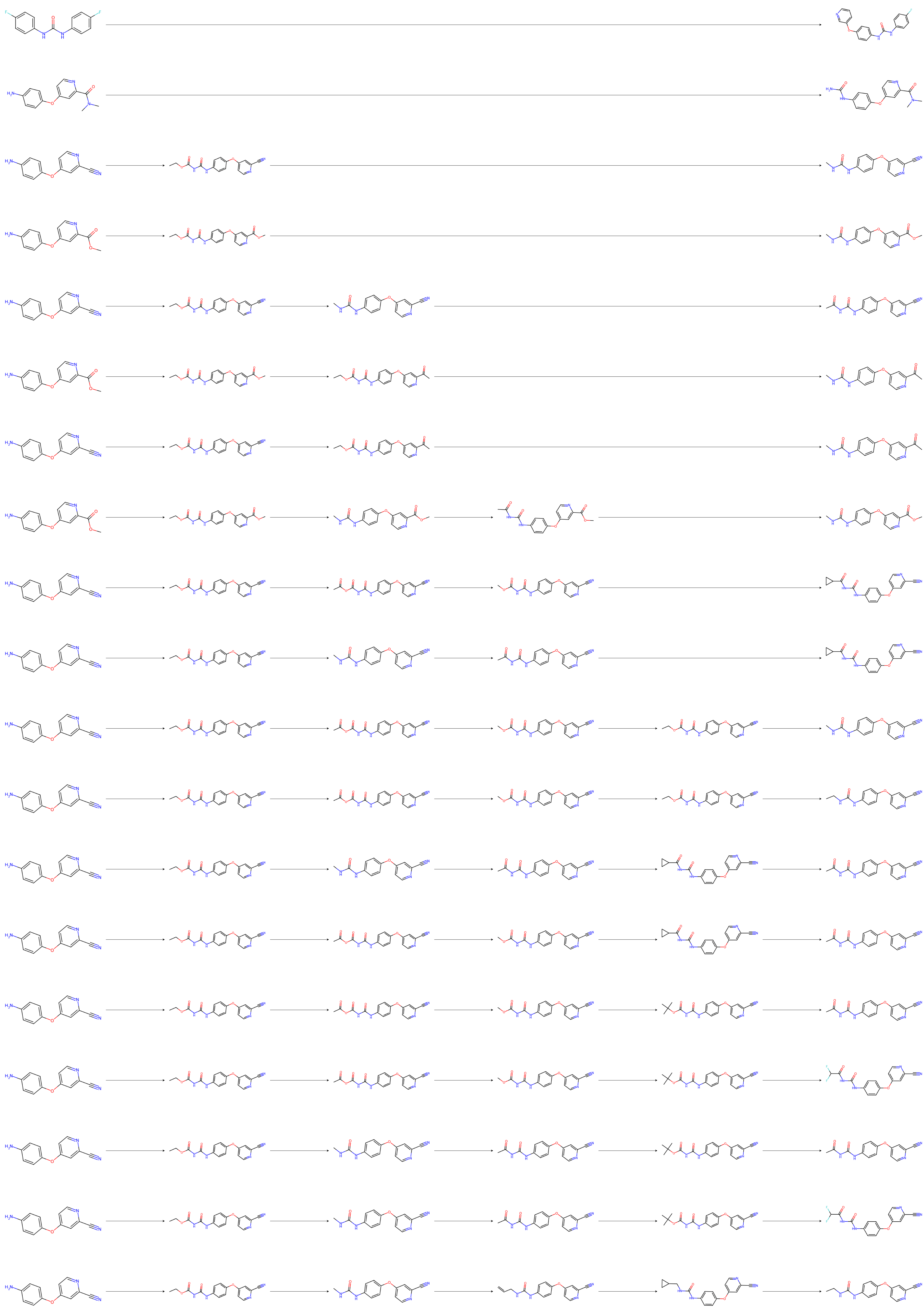

P1394
P455
P1689
P1173
P1070
P687
P687
P1173
P248
P248
P1689
P1573
P1070
P1070
P1070
P1692
P1070
P1692
P1573
